# A protocol for lab-scale production of ^13^C yeast extract as internal standard for metabolomics and quantification of intracellular metabolites

**DOI:** 10.64898/2026.06.12.731807

**Authors:** Miriam Cammaert, Rosalie Wouters, Jitske van Ede, Erik de Hulster, Christiaan Mooiman, Patricia van Dam, Martin Pabst, Walter van Gulik, Pascale Daran-Lapujade

**Author notes:** These authors contributed equally to this work (Miriam Cammaert, Rosalie Wouters, Jitske van Ede).

## Abstract

Metabolomics enables the profiling of small-molecule metabolites and thereby captures the biochemical state of a living organism at a given moment and enables to monitor its cellular responses to stimuli. This technique has become a powerful tool in pharmaceutical research, the food industry, and microbial research. Metabolomics aims to obtain an unbiased metabolic profile; however, this is complicated by compound instability, complex and often extensive sample processing, and nonlinear responses in mass spectrometry. Therefore, correcting for metabolite loss and mass spectrometry-related artifacts is essential, typically achieved through relative quantification against an isotopically labelled internal standard for each metabolite of interest. This article describes how to produce ^13^C-labelled yeast extract and its use as internal standard for metabolomics. More specifically, it provides step-by-step protocols for the fed-batch fermentation, quenching, metabolite extraction, and LC-MS and GC-MS characterization of the internal standard. It also includes a protocol explaining how to use the internal standard for the quantification of metabolites in yeast samples.

## Introduction

Metabolic profiling is a powerful tool for characterizing biological systems. By detecting and quantifying many intracellular metabolites simultaneously in a biological sample, metabolomics captures the biochemical state of a living organism at a given moment and enables to monitor its cellular responses to stimuli. Metabolomics is an established technique that is used in both fundamental and applied research in various fields, including pharmaceutical research (Wishart, 2016), the food industry (Li et al., 2021), and microbial research (Sailwal et al., 2020; Tang, 2011).

The aim of metabolomics is to obtain an unbiased *in vivo* snapshot of the metabolic profile of a living organism. This aim is however difficult to achieve as metabolites can be unstable and lost during the extensive processing required to quench metabolic reactions and to extract metabolites from cells and tissues (Canelas et al., 2009). Furthermore, differences in sample matrices between organisms can cause variation in ionization efficiency and signal suppression or enhancement in mass spectrometry (Panuwet et al., 2016). Finally, mass-spectrometry-based quantification of metabolite levels is error-prone as high salt or organic compound concentrations in the sample can lead to ion suppression in the electrospray ionization step, causing a nonlinear response of the metabolite levels (Gowda & Djukovic, 2014). These technical challenges can be mitigated by using a method known as isotope dilution mass spectrometry (ID-MS) (Mashego et al., 2004; Wu et al., 2005). The principle of ID-MS is based on the addition of a stable isotopically labelled internal standard for every metabolite of interest. ID-MS thereby allows for quantification relative to the internal standard rather than absolute quantification, correcting for metabolite loss and the effects of ion suppression.

Internal standards - typically ^13^C or ^15^N isotopically labelled metabolites - must meet several criteria. First, the internal standard should be uniformly isotopically labelled. Second, the internal standard should mimic the metabolite profile of the sample to maximize the number of quantifiable metabolites. Third, the internal standard should be readily available and cost-effective. Finally, the internal standard should contain metabolites at concentrations closely matching those in the target sample to ensure accurate quantification.

These internal standards can be commercially available or ‘home-made’. Although commercially available internal standards offer the benefit of quality control, they are at risk of discontinuation. Furthermore, commercial internal standards are generic and not optimized for the organism and/or condition of interest, which may hinder the accurate measurement of crucial metabolites. These caveats can be circumvented by producing in-house isotopically labelled internal standard.

The in-house production of internal standards requires several decisions aimed at optimizing the quality of the standard in terms of metabolite diversity, abundance, and preservation during processing. These choices include the organism (Seike et al., 2025) and the type of (labelled) substrate depending on the metabolic profile of interest. Furthermore, the type of culture (e.g., batch, fed-batch, or chemostat) should minimize ^12^C contamination and maximize biomass yield on substrate. The harvesting and extraction procedure should be chosen as to minimize degradation and leakage of metabolites. Finally, the choice of analytical methods should take into consideration the metabolite chemical properties and their concentration range (Figure 1).

**Figure 1.**
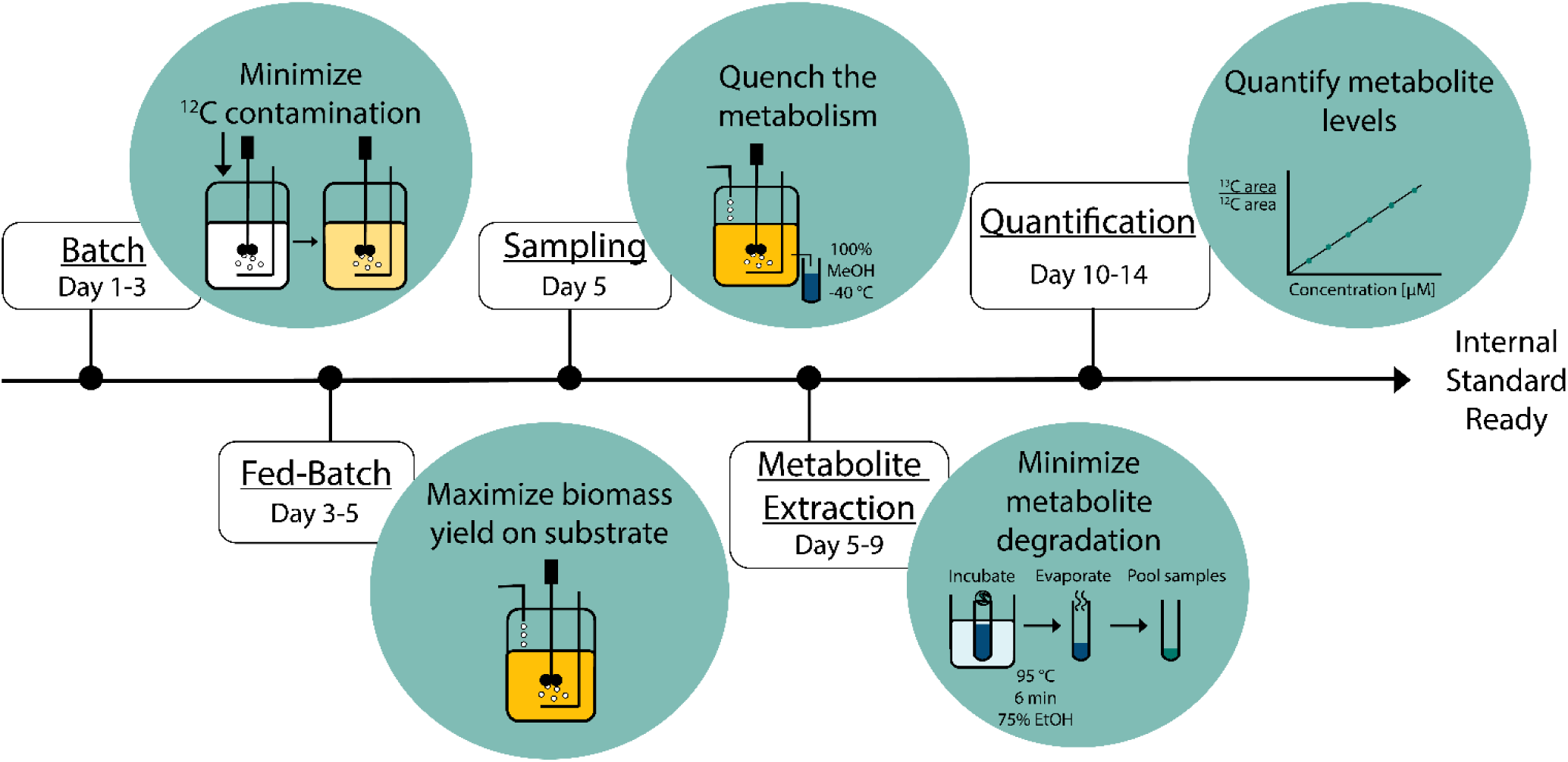
Timeline for in-house production of ^13^C-labelled internal standard.

The present study describes a robust protocol for the in-house production of internal standards, based on fully ^13^C-labelled yeast biomass. The protocol outlines cultivating fully labelled *Saccharomyces cerevisiae* (*S. cerevisiae*) in a high-density fed-batch, rapid harvesting and quenching of the biomass, methanol-based metabolite extraction, and quantification of metabolites in the resulting internal standard using liquid-chromatography and gas chromatography coupled to mass spectrometry (abbreviated as LC-MS and GC-MS, respectively). The final protocol describes how to use internal standards to quantify intracellular intermediates in NAD^+^ biosynthesis. While this method focuses on *S*. *cerevisiae*, the described internal standard is relevant for all types of organisms, particularly for the most commonly investigated metabolic pathways (e.g., within the Embden-Meyerhof-Parnas (EMP) pathway of glycolysis, the tricarboxylic acid (TCA) cycle, the pentose phosphate pathway, etc.). Furthermore, metabolites absent in *S. cerevisiae* can be added in labelled form to the internal standard if desired. The presented workflow highlights key aspects for the production of internal standards that can be customized to suit different experimental conditions, lab capacities, and microbes.

### Strategic planning

Note that this protocol requires basic knowledge in microbial physiology and fermentation technology (Heijnen, 2009) and access to an analytical chemistry facility. The production process of ^13^C-labelled internal standards must meet two main requirements: ensure full isotopic labelling of the microbial biomass and maximize the biomass yield on substrate to minimize production costs.

To obtain fully labelled biomass, all non-labelled sources that might contaminate the metabolites must be removed, while keeping a reasonable growth rate. For *S. cerevisiae*, vitamins whose carbon atoms are incorporated into metabolites are omitted from the medium. Removal of pantothenic acid, nicotinic acid, myo-inositol, pyridoxine, and 4-aminobenzoic acid from the medium decreases yeast specific growth rate by a factor of six. In contrast, biotin and thiamine, which act exclusively as cofactors without donating carbon to metabolites, were supplemented. For vitamins that cannot be omitted but might end up in carbon, we advise adding commercial ^13^C-labelled versions. The same principle applies to other potentially essential nutrients, such as amino acids. Another measure to limit both the contamination by unlabelled carbon and loss of substrate, is to inoculate the batch fermentation with a very small amount of unlabelled biomass. Typically, bioreactors are inoculated at 1-5% (v/v) of the working volume using a preculture of the same cultivation condition. However, preparing precultures with labelled glucose would result in the loss of a significant amount of costly ^13^C-labelled glucose. The bioreactors can be inoculated with unlabelled biomass provided it represents a minute fraction of the final total biomass. This is achieved by inoculating the batch cultures with a single 1 mL cryo-vial of unlabelled *S. cerevisiae*.

To maximize the biomass yield on substrate, it is preferable to favour the most efficient metabolic mode in terms of ATP yield on substrate, which is typically respiration. Consequently, fermentative substrate dissimilation should be prevented or minimized. For instance, *S. cerevisiae* is a Crabtree-positive yeast that ferments glucose when supplied in excess, even in the presence of excess oxygen, leading to decreased biomass yield on carbon substrate (Y_x/s_) as compared to fully respiratory growth (Y_x/s_ respiration of 0.5 g_biomass_/g_glucose_ vs Y_x/s_ fermentation of 0.1 g_biomass_/g_glucose_) (De Deken, 1966; Van Dijken et al., 2000). To ensure fully respiratory growth, aerobic glucose limitation is required, which can be reached using glucose-limited fed-batch cultures. Conversely, Crabtree-negative yeasts do not require glucose-limited conditions and batch fermentations with excess carbon source can be used, provided oxygen is sufficient to prevent oxygen limitation. It is also important to choose the growth conditions, medium, and gas supply for production of the internal standards that mimic as closely as possible the conditions in which the experiments will be performed. For instance, internal standards produced with glucose as carbon source will not contain labelled intermediates of the Leloir pathway that are only produced when galactose is used as carbon source.

Upon harvesting, the extracellular concentrations of glucose and oxygen—key regulators of metabolism— are taken up by the cells and depleted within milliseconds in the samples (Koning & Dam, 1992), leading to rapid alterations in intracellular metabolite levels, particularly those with turnover times in the order of seconds. As a result, any delay between harvesting and quenching may compromise the metabolic profile. Therefore, a fermentor equipped with a rapid bottom-harvesting port is necessary to enable rapid harvesting (Lange et al., 2001). Given the sensitivity of glucose-limited cultures to glucose and oxygen perturbations, rapid harvesting combined with instantaneous quenching is essential to ensure reliable metabolic profiling. Quenching is an organism-dependent method that requires careful optimization to prevent leakage of metabolites in the extracellular matrix and their degradation. Instantaneous quenching of various yeast species, including *S. cerevisiae*, *Kluyveromyces marxianus*, and *Pichia pastoris* (recently renamed *Komagataella phaffii* (Bolten & Wittmann, 2008)), can be achieved using -40 °C methanol (Canelas et al., 2008).

After quenching the cells in ice-cold methanol, the samples are further processed, first to separate intracellular from extracellular metabolites. Subsequently, metabolite extraction is performed using boiling ethanol (Canelas et al., 2009). After extraction, ethanol is removed by vacuum evaporation. The resulting residues cell extract is resuspended in water, filtered, and subsequently processed either by solid-phase extraction (SPE) or by derivatization prior to metabolomic analysis using LC–MS or GC–MS, respectively.

The basic protocols are separated into **trial run** and **real run**. A trial run is strongly recommended to evaluate the microbial growth, biomass concentration, and physiology under the modified culture conditions. The trial run should be conducted using ^12^C-glucose, which significantly reduces costs and permits sampling in time to assess the fermentation profile. The final biomass concentration achieved during the batch phase is critical data to define the optimal glucose feeding rate for the fed-batch phase. Overfeeding will result in respiro-fermentative metabolism and loss of ^12^C-glucose in the form of ethanol, glycerol and organic acids, while underfeeding leads to slow growth rate and associated decrease in biomass yield on substrate due to maintenance requirements (see Support Protocol 1) (Verduyn, 1991). Additionally, the trial run provides the perfect opportunity to practice the rapid harvesting protocol, and the subsequent sample processing and metabolite extraction. Due to the sensitivity of metabolic profiles to environmental changes, teamwork and a streamlined workflow are essential. Following the fermentation and sample processing, at least one unlabelled metabolite sample should be analysed to confirm that the metabolite profile of interest is obtained under the tested conditions. These validation steps form the foundation for a successful ^13^C-labelling experiment, during which intermediate sampling is typically avoided due to the high cost of isotopically labelled substrates.

This protocol first describes how to perform the trial run for the fed-batch (Basic Protocol 1), followed by the sample processing and metabolite extraction (Basic Protocol 2), and the verification of the metabolite profile of this trial run (Basic Protocols 3 and 4). The protocol then describes how to perform the fed-batch with ^13^C-labelled glucose (Basic Protocol 5) and how to characterize the internal standard (Basic Protocols 6 and 7). The last protocol (Basic Protocol 8) describes how to quantify metabolites in yeast samples from shake flask cultures using the internal standard.

*NOTE*: To allow handling of the samples as quickly as possible for harvesting and sample processing (Basic Protocols 1 and 2), gathering a team of at least six people for the harvesting day is convenient to distribute the tasks. Two persons are responsible for the harvesting, one person adjusts the feeding profile, two persons are responsible for the sample processing, and one person is responsible for changing the centrifugation rotors.

## Basic Protocol 1: Trial Run – Fed-Batch

It is strongly recommended to perform trial run(s) with unlabelled glucose. During the trial run, you will prepare and execute an aerobic batch and fed-batch fermentation, validate the fed-batch feeding profile, test the rapid harvesting set-up and produce samples for optimization of the analytical pipeline. This protocol describes how to perform an aerobic batch and fed-batch fermentation in a bioreactor. The growth performance of *S. cerevisiae* is monitored via sampling for biomass and extracellular metabolites and via continuous gas analysis. This data will inform whether the culture conditions are optimal for a high biomass yield on substrate. Finally, biomass will be harvested from the bioreactors in large quantities using a rapid harvesting device, giving the opportunity to further optimize the rapid harvesting routine if needed. Once this protocol has been successfully performed, you are ready to continue to Basic Protocol 2: Quenching and washing of the cell samples and metabolite extraction.

### Materials

#### Media and chemicals

*S. cerevisiae* strain CEN.PK113-7D or other strain (1 mL in 30% v/v glycerol)

Vitamin solution (see recipe in Reagents and Solutions)

Trace elements solution (see recipe in Reagents and Solutions)

Batch salt base (see recipe in Reagents and Solutions)

Fed-batch medium (see recipe in Reagents and Solutions)

12C-Glucose solution (see recipe in Reagents and Solutions)

Antifoam C solution (see recipe in Reagents and Solutions)

Methanol (HPLC-grade >99.9%) (-40 °C)

pH calibration solution, 4 and 7

Base solution (4 M NH_4_OH, see recipe in Reagents and Solutions)

Sterilized demi-water, 50 mL

### Bioreactor equipment

Fully equipped fermentor with a working volume up to 1.25L, including bottom rapid harvesting port and

several pumps, connected to ez-Control (e.g. Applikon, Getinge, Delft, the Netherlands)

Norprene (Masterflex) tubing

At least 3 screwcaps for Schott bottles with two in/outlets

Glass shake-flask with outlet tube at the bottom

pH probe (sterilizable, and calibrated between a pH of 4 and 7)

DO probe (sterilizable)

Platinum temperature sensor

Pump (0.02-0.5 mL/min range, for fed-batch feed)

Fixed speed pump (20 rpm with Norprene 13 or 14 tubing allowing for 1.2 – 5.2 mL/min) (for antifoam and base)

Pressure controller (able to reach 0.3 bar overpressure on the bioreactor) NGA 2000 Rosemount gas analyser (e.g. Emerson, St. Louis, MO)

Scale connected to the control system to log the decrease of fed-batch medium in the bottle (up to 6200 grams)

#### Harvesting equipment

Schott bottles (1L)

Schott bottles filled with 450 mL methanol, a stirrer, and with a volume indicator at 550 mL (or 575 mL depending on the harvesting schedule)

Scale to measure the volume of harvested culture (up to 6200 grams)

Magnetic stirrer

Stirrer bar

Cryostat with cryofluid (70% v/v ethylene glycol), set at -40 °C

Freezer, set at -40 °C

Centrifuge (tabletop, max. speed 17000 x g)

#### General lab equipment

Sterile syringe (up to 2 mL)

Syringe filter (0.22 µm pore size)

HPLC vials

Agilent 1260 HPLC, equipped with a Bio-Rad HPX 87H column Equipment for dry weight determination (see Support Protocol 1)

Cryo and nitrile gloves

*CAUTION:* NH_4_OH has a high vapour pressure, and the gas is toxic. Always work in a fume hood, and **do not** sterilize this solution. Rather, autoclave a bottle with water and add NH_4_OH in a fume hood afterwards.

*CAUTION:* When working with high molarity acids and bases, please take the required precautions like wearing protective clothing, glasses and gloves.

*CAUTION:* Methanol is a neurotoxic compound and can penetrate the skin. Always wear (nitrile) gloves when handling methanol and work as much as possible in the fume hood. Try to minimize the steps performed outside the fume hood.

*CAUTION*: The fermentor is operated under pressure at 0.3 bar. Do not open any ports and/or valves without first depressurizing the reactor. Sudden release of pressure may cause injury or equipment damage.

*CAUTION*: Ensure that the gas analyser used for off-gas CO₂ monitoring can detect isotopically labelled CO₂. Infrared (IR)-based sensors may not accurately measure ¹³CO₂ because its absorption spectrum is shifted relative to ¹²CO₂ due to the higher isotopic mass. Using an analyser calibrated only for ¹²CO₂ can therefore lead to significant underestimation or misinterpretation of CO₂ levels.

### Bioreactor preparation

1. (Deep) Clean a Rapid-Harvesting Fermentor (2 L) and replace all Norprene (Masterflex) tubing (Figure 2).
2. Calibrate the pump for fed-batch media inflow (flow range for 0.02 to 0.5 mL/min).
3. Prepare batch medium without vitamins and glucose and add the 350 mL batch salt base to the fermentor.
4. Calibrate the pH probe (Figure 2, number 7).
5. Sterilize the fermentor at 121 °C and let it cool down.
6. Calibrate the DO probe (Figure 2, number 6).
7. Attach the base (or acid; Figure 2, number 13) and antifoam bottles (with screwcaps with two or three in/outlets; Figure 2, number 15) to the bioreactor, and ensure tubing is already filled with fluid before the fermentation starts.
8. Set temperature control to 30 °C and pH control to 5.

### Protocol

1. Set the gas inflow to 0.5 L/min and the stirrer speed to 500 rpm.

**Figure 2.**
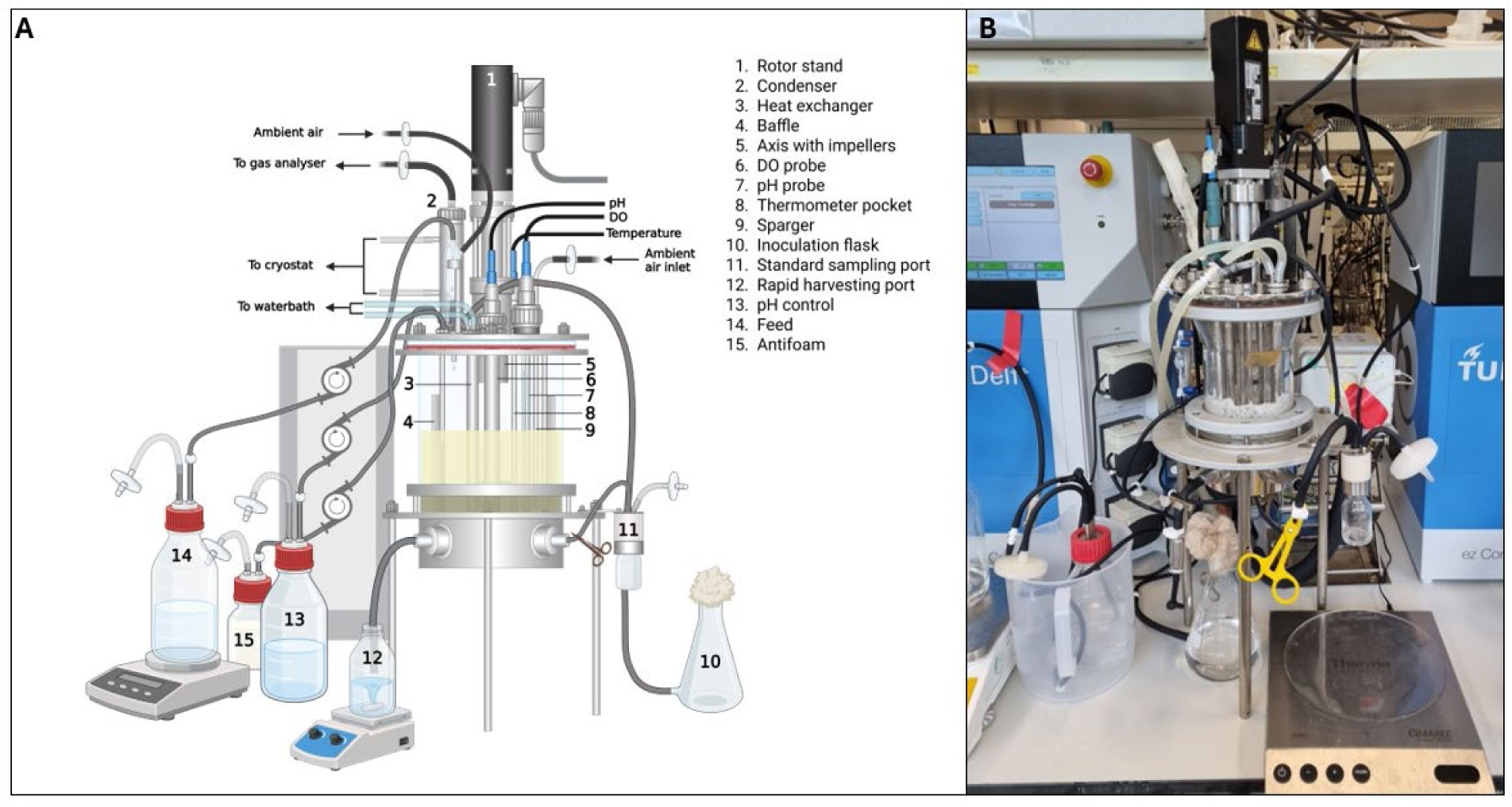
Schematic overview and photo of the fermentation and harvesting set-up. A. Schematic representation of the fed-batch and rapid harvesting set-up. Created in https://BioRender.com. B. Picture of the fed-batch set-up at the end of the fermentation.

2. Add 100 mL glucose solution (glucose starting concentration of 75 mM), 1.5 mL filter sterilized vitamin solution, and 1 mL CEN.PK113-7D glycerol stock to the inoculation flask (Figure 2, number 10) and add to the fermentor.

3. Flush the inoculation flask by adding 50 mL sterilized demi-water and transferring it to the fermentor.

The starting volume in the bioreactor is ca. 500 mL.

4. Set the pressure on the bioreactor to 0.3 bar.

5. In case of excessive foaming, add a few drops of antifoam by briefly switching on the pump.

6. Sample at regular time intervals for extracellular metabolites and dry weight measurements using a standard sampling port (Figure 2, number 11). See Support protocol 1: Dry weight determination for the procedure for dry weight measurements.

7. Continuously monitor the CO_2_ profile over time using a gas analyser. Arrest in CO_2_ and biomass production indicates that all carbon is depleted in the batch phase and that the fed-batch can be started (Figure 3).

**Figure 3.**
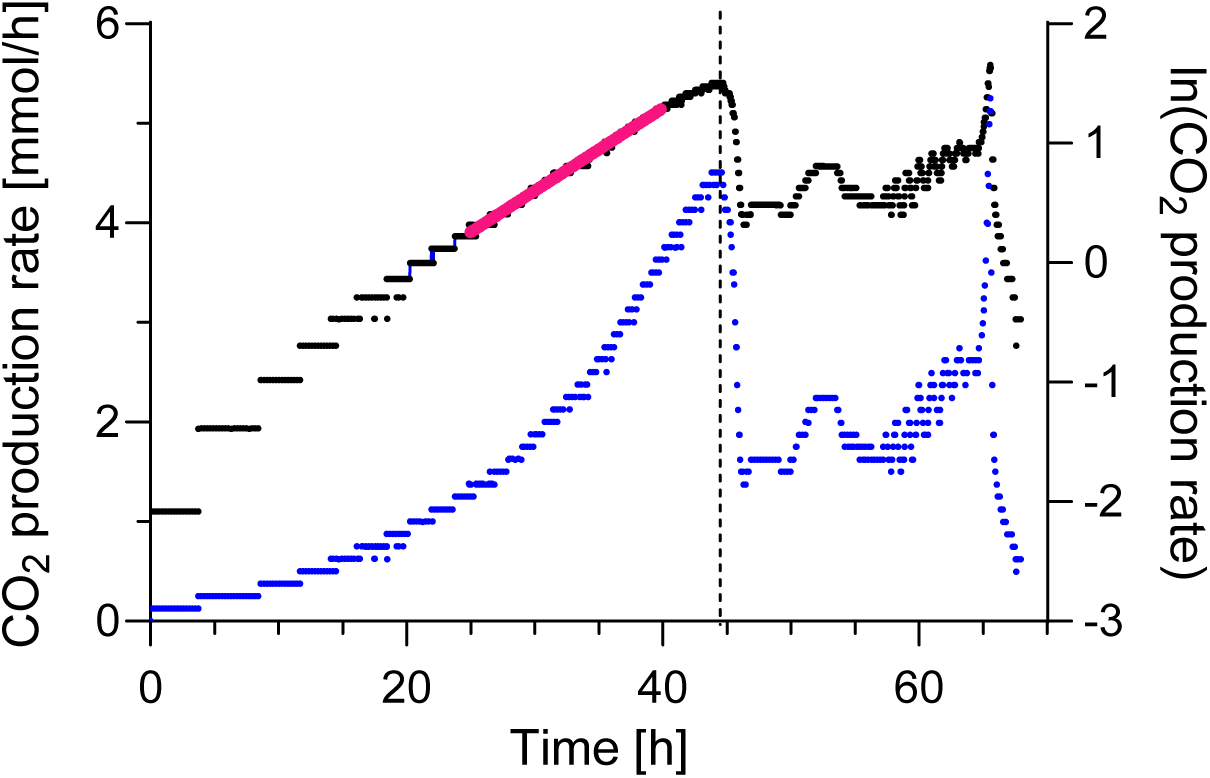
CO2 production profile during the batch phase. The CO2 production rate [mmol/h] is depicted in blue (left y-axis) and the natural logarithm (ln) of the CO2 production rate is depicted in black (right y-axis). The trendline through the ln(CO2 production rate) gives an approximation of the maximum growth rate. For this culture, using timepoints between 20 hours till 40 hours (purple line) gives an estimated growth rate of 0.0692 h^−1^ (linear regression equation y = 0.0692x - 1.4735 with R² = 0.995).

8. Calculate the fed-batch feeding profile based on the final biomass concentration after the batch fermentation following equations 1 and 2 (Heijnen, 2009; Hensing et al., 1994).

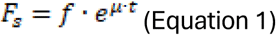

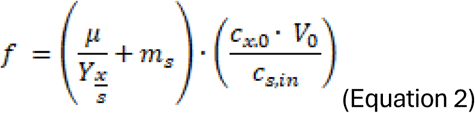

With:

F_s_ [L h^−1^] = volumetric substrate feed rate

f [L h^−1^] = feed factor

t [h] = time

µ [h^−1^] = growth rate

Y_x/s_ [g_s_ g_x_^−1^] = yield of biomass (x) on substrate (s), organism-dependent

m_s_ [g_s_ g_x_^−1^ h^−1^] = maintenance requirements, organism-dependent

c_x,0_ [g_x_ L^−1^] = biomass concentration in the bioreactor at the end of the batch phase

V_0_ [L] = volume in the bioreactor at the end of the batch phase

c_s_,in [g_s_ L^−1^] = substrate concentration in the feed

*The Y_x/s_ and m_s_ values are dependent on the organism and the cultivation conditions. For* S. cerevisiae *grown on glucose, the Y_x/s_ and m_s_ are 0.5 g_s_ g_x_^−1^ (van Hoek et al., 2000) and 0.0072 g_s_ g_x_^−1^ h^−1^ (Vos et al., 2016), respectively. These values can be found in literature for various model organisms or can be experimentally determined using chemostat cultures. In the fed-batch phase the specific growth rate, µ, is set by the feeding rate. If the specific growth rate is set to a value above the maximum growth rate of the organism, the feeding rate will be too high and glucose will be supplied in excess, resulting in respiro-fermentative metabolism. It is therefore important to determine the maximum specific growth rate of the organism under the chosen cultivation conditions, for instance by measuring biomass concentration (optical density) during the batch phase or by performing a separate shake flask experiment. For* S. cerevisiae *under the given cultivation conditions, the maximum growth rate measured in batch culture is approximately 0.07 h^−1^ (Figure 3) and the growth rate for the fed-batch phase was set at 0.05 h^−1^. The initial volume in the bioreactor at the start of the fed-batch is 0.5 L and the substrate concentration in the feed is 116.41 g L^−1^. Depending on the total working volume of the bioreactor, sparging and gas inflow capacity, and feeding pump capacity, the initial volume in the bioreactor and substrate concentration of the feed might have to be adjusted. Finally, to calculate the feeding profile, the biomass concentration must be accurately measured (see Support protocol 1: Dry weight determination)*.

9. Start the fed-batch by switching on the feeding pump with fed-batch medium at the calculated rate.

*Verify the unit of the flow rate required by the pump controller—whether it is specified in mL/min or L/h. Additionally, confirm whether the exponential factor - the e^µt^ (Equation 1) is applied automatically by the software, or if it needs to be manually adjusted within the program*.

10. Take samples for extracellular metabolites determination and dry weight determination (see Support protocol 1: Dry weight determination) at regular intervals during the fed-batch fermentation to monitor the progress and to check whether the biomass concentration is in line with the expected profile, see Supplementary File 1 for the simulated biomass profile during the fed-batch (Figure 4). For extracellular metabolites, spin down samples, transfer supernatant to a vial and analyse substrate (here glucose) and product (ethanol and organic acids for yeast) concentrations by standard gas or liquid chromatograph methods.

**Figure 4.**
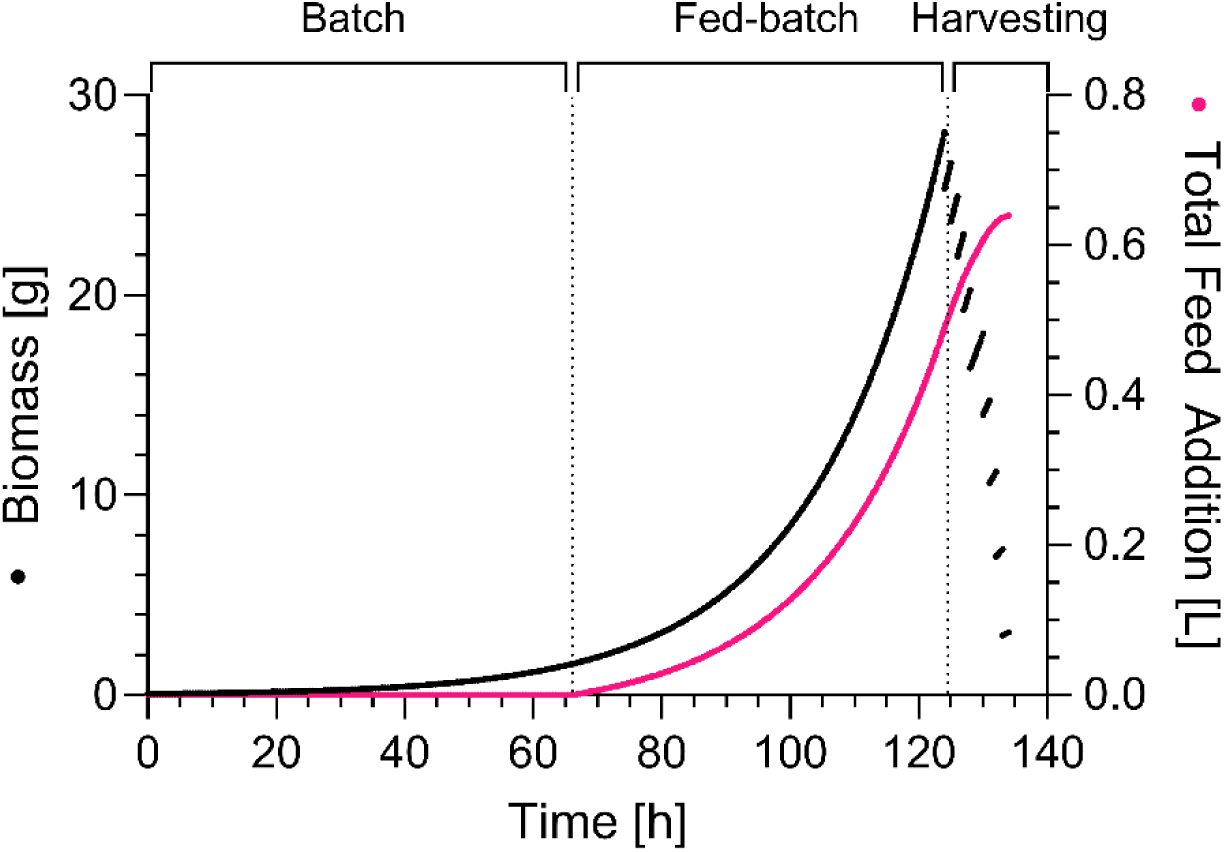
Simulated medium feeding and biomass profiles during the fed-batch and harvesting phases. The cumulative volume of medium fed to the bioreactor (right axis) and corresponding biomass profile (left axis) were extrapolated from the dry weight determination at the end of the batch phase.

*We recommend taking at least five samples during the fed-batch phase to ensure that the feeding profile has been set correctly*.

11. Harvest at regular intervals from the bioreactor.

*The harvesting will mimic the harvesting phase that will be performed at the end of the real run with ^13^C-glucose and is a good rehearsal for the harvesting team. The harvesting time should be carefully chosen based on two factors. On one hand, it is important to perform the harvesting phase while medium is still supplied to the culture to prevent carbon starvation and depletion of metabolites. On the other hand, as little as possible of the ^13^C-glucose containing medium should be left unused. To prevent excess feeding of glucose during the harvesting phase, the feeding rate will be adjusted according to the broth volume remaining in the fermentation, see step 16. When sampling starts, attach two -40 °C 50 mL rotors to the centrifuges*.

*To check when the medium will be exhausted look at cell 4E in Supplementary File 1, spreadsheet “FEDBATCH_SAMP”. The volume in the bioreactor after harvesting should be as close to zero as possible, see cell 58E in the same sheet*.

12. Take a Schott bottle with 450 mL of 100% methanol from the freezer (-40 °C) immediately before harvesting, weigh rapidly and write down the weight. This step is important to monitor how much volume is removed from the fermentor.

13. Put the bottle under the rapid harvesting port (Figure 2, number 12) on a stirring plate and turn on the stirrer. Turn on the stirrer at a steady, medium speed so that the broth and methanol mix as quickly as possible while minimizing the risk of methanol splashing.

14. Sample approximately 100 mL of broth directly in the Schott bottle with methanol. Use the volume indicator on the bottle at 550 mL as a guideline.

*The volume of harvested broth is limited by the capacity to process samples rapidly. For instance, the number of tubes that can be spun down simultaneously, the number of people available for fast handling of the samples, but also the -40°C freezer capacity. If the centrifugation capacity in the lab is limited to six 50 mL Greiner tubes per round, the additional six 50 mL Greiner tubes can be stored in a -40 °C freezer to avoid the samples from heating up too much. A critical requirement for proper quenching is that the harvesting is always performed in a 1:5 broth to methanol ratio (Mashego et al., 2004)*.

15. Weigh the bottle with broth/methanol mixture and calculate the sampled volume.

16. Adjust the feed rate according to the harvested broth to keep the growth rate constant. Put the recorded weight of the broth in cell O17 for the first sample, and O22 for the second sample, and so on in Supplementary File 1, spreadsheet “FEDBATCH_SAMP”. Based on the sampling volume, the feed rate is automatically calculated, see cell J18 for the adjusted feed rate in L/h and cell K18 for the feed rate in mL/min.

*For a detailed explanation of the calculations check Supplementary File 2. Briefly, first calculate the new volume in the bioreactor, based on V = V0 + V_feed – V_sampling. In this formula, V_feed is obtained by integration of equation 2; ΔV_feed = ∫ Fs = (f/mu)*exp(mu*t) - (f/mu), and V_sampling is obtained by weighing the sample. Next, adjust the total amount of biomass in the reactor after harvesting (M_X_#) which can be determined based on the volume of the sample (ΔV_sample) as a fraction of the total volume (V); M_X_# = M_X * (1 - (ΔV_sample/V)). Based on the adjusted volume and biomass concentration, the adjusted feed rate can be calculated according to equation 2*.

*Due to the discrepancy between exponential increase of the total volume in the reactor and the total amount of biomass in the reactor, it is important to calculate the total amount of biomass rather than the biomass concentration for determining the feeding profile*.

17. Proceed with Basic Protocol 2 for metabolite extraction.

18. Repeat steps 11-17 until the entire bioreactor is empty.

S. cerevisiae *cultures with fully respiratory metabolism on glucose have a respiratory quotient (RQ, ratio of the carbon dioxide production rate to the oxygen consumption rate), around 1.06. Respiro-fermentative metabolism will lead to increased RQ. On-line monitoring of the culture RQ therefore enables to assess whether the feed rate is set in the correct range. If the RQ exceeds 1.2, it is advised to adjust the feed rate accordingly. Figure 5 depicts the RQ profile during batch and fed-batch culture of* S. cerevisiae*. The RQ is high (above 2) during the glucose consumption phase in batch, then decreases around 0.5 during the following ethanol and organic acids consumption phase in batch and subsequently stabilizes around 1.1 in the fed-batch phase*.

**Figure 5.**
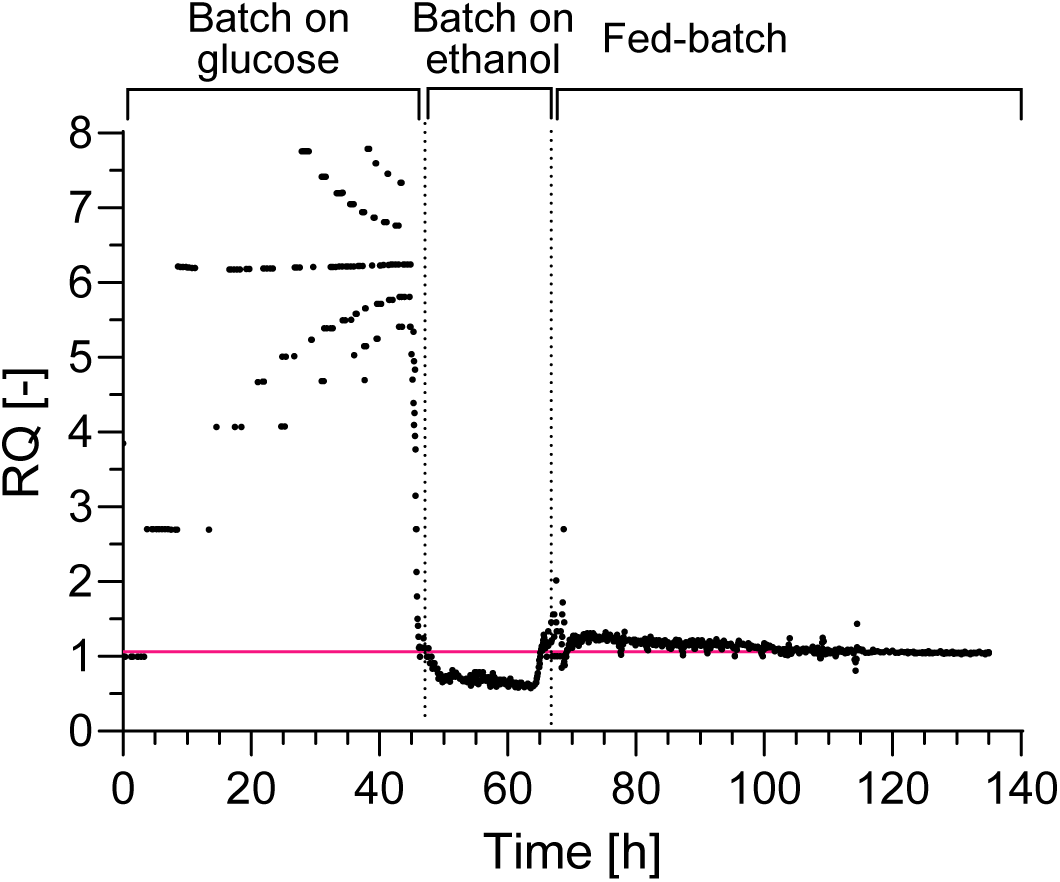
Respiratory Quotient profile during the fermentation. The Respiratory Quotient (RQ), ratio of the carbon dioxide production rate to the oxygen consumption rate, is used as proxi for the metabolic mode during fermentation. The pink line indicates fully respiratory metabolism with an RQ of 1.06. The scattering of RQ values in the batch phase reflects the low biomass concentration in the bioreactor and resulting low CO2 production and O2 consumption rates that approach the detection limit of the gas analyser.

## Support protocol 1: Dry weight determination

This protocol describes how to determine the dry weight of the culture, according to Postma et al. (1989). Always perform the dry weight determinations in duplicate.

### Materials

Demi-water Culture sample

Nitrocellulose filters (pore size 0.45 µm)

Microwave Oven (80 °C) Glass Petri dish Flat tip tweezer

Precision balance (>=0.001 g)

Vacuum-pump connected to filtering device

Calibrated volumetric pipette (10 mL)

### Protocol

1. Dry the filters, for a minimum period of 8 hours at 80 °C in an oven. The filters can be kept in the oven for prolonged periods of time.
2. Place the blue separation paper disk included in the filter box in a dry glass Petri dish. Collect the dry weight filter, place it on the blue disk and close the petri dish. *Always handle the filter using a flat tip tweezer, avoid bringing the filter in contact with moist materials*.
3. Weigh the dried filter, without the blue separation paper, on a precision balance, and wait until the signal stabilises.
4. Switch on the vacuum-pump connected to the filtering device and place the dried filter on the filtering device.
5. Wet the filter with approximately 10 mL of demi-water.
6. Mix the culture sample, by shaking, and pipette with a calibrated volumetric pipette exactly 10 mL of the culture sample slowly on the filter. For cultures with low biomass concentration (below 0.3 g/L), adjust the volume of sample (e.g. 20 mL).
7. Carefully wash the filter with the cell broth with 20 mL of demi-water to remove salts. Avoid splattering.
8. Remove the filter from the filtering device using the tweezer (be careful not to touch the cell material).
9. Place the blue separation disk on the microwave dish and place the filter on top of the disk to prevent sticking of the filter to the microwave dish.
10. Dry the filter in the microwave at low power settings, e.g. 350 W for 20 minutes using defrost.
11. Immediately after drying, weigh the filter on the precision balance.
12. Calculate the dry weight in grams per litre by subtracting filter mass from filter mass with biomass. Deviation between duplicates should be lower than 1%.

## Basic Protocol 2: Quenching and washing of the cell samples and metabolite extraction

This protocol describes how to rapidly quench cellular metabolism by direct mixing of the broth into cold methanol, prevent contamination of cell extracts by metabolites present in the medium by cold centrifugation and washing, and how to extract the metabolites from the cells with boiling ethanol. After metabolite extraction, ethanol is evaporated, cells are resuspended in water, and the resulting metabolite extract is separated from cell material by centrifugation and filtration. The same protocol is used to produce cell extracts from the trial run (^12^C only, Basic Protocol 1), from the real run (^13^C only, Basic Protocol 5), and to finally quantify metabolite concentrations in samples of interest using internal standard (^12^C extract with ^13^C-labelled internal standard, Basic Protocol 8). For the latter, internal standard is spiked before the metabolite extraction using boiling ethanol at step 25. Upon successful completion of this protocol, the cell extract is aliquoted and stored at – 80 °C.

### Materials

Methanol (HPLC-grade >99.9%), -40 °C

Methanol (83% v/v with demi-water), -40 °C

Ethanol (75% v/v) (>99.8%)

MilliQ

Freezer, -40 °C (for methanol storage), -80 °C (for sample storage)

Cryostat with cryofluid, capable of reaching -40 °C

Centrifuge (tabletop, -19 °C); one rotor for 15 mL tubes and one rotor for 50 mL tubes (stored at -40 °C)

Two bottle-top dispensers (5 mL and 20 mL capacity)

Waste bucket for methanol-contaminated waste (plastic and liquid)

Two water baths (75 °C, and 95 °C)

Bucket with ice (to fit 12 x 50 mL Falcon tubes comfortably)

Regular marbles (14 mm diameter) Falcon tubes (50 mL, 135 pieces)

Sampling tubes (15 mL, 135 pieces; need to fit the SpeedVac inlets) Schott bottles (1 L)

Vortexes

Benchtop centrifugal vacuum concentrator (SpeedVac), e.g. Labconco CentriVap or equivalent, with vacuum pump and cold trap, capable of evaporating ethanol and equipped with a rotor for 15 mL conical tubes

Syringes

Syringe filters (0.2 μm pore size, PVDF)

Nitrile gloves

#### Protocol

1. Turn on the cryostat(s), and centrifuge(s) to reach the desired temperature.
2. Put all four bottles filled with 83% (v/v) methanol solution in the cryostat and attach a bottle-top dispenser to one of the bottles.
3. Turn on the water baths at 75 °C and 95 °C.
4. Put 12x 15 mL Falcon tubes containing 75% (v/v) ethanol, or a Schott bottle filled with 75% ethanol and equipped with a bottle-top dispenser, in the 75 °C water bath. *Always keep the samples on ice, at least until the boiling ethanol step (step 26)*.
5. Harvest the broth in cold methanol following the instructions of Basic Protocol 1.
6. Immediately after harvesting the broth into the cold methanol, transfer the quenched broth to 12x 50 mL falcon tubes (appr. 48 mL per tube) by pouring. *Pouring of the broth-methanol solution should be executed in a fume hood for safety reasons*.
7. Centrifuge (5 min, 2500 x g, -19 °C).
8. Decant the supernatant into the methanol waste vessel.
9. Add 20 mL of 83% (v/v) methanol (-40 °C) to all pellets using the bottle-top dispenser.
10. Resuspend the pellets by vortexing.
11. Centrifuge (5 min, 2500 x g, -19 °C).
12. Change the rotor for 50 mL tubes to the rotor for 15 mL tubes and put the other rotor back at -40 °C.
13. Adjust the bottle-top dispenser on the 83% (v/v) methanol to 3 mL.
14. Decant the supernatant into the methanol waste vessel.
15. Add 3 mL of 83% (v/v) methanol (-40 °C) to all pellets using the bottle-top dispenser.
16. Resuspend the pellets by vortexing.
17. Pour the liquid into the 12x 15 mL sampling tubes.
18. Wash the 12x 50 ml Falcon tubes with 3 mL 83% (v/v) methanol (-40 °C) using the bottle-top dispenser.
19. Pour the liquid to the 12x 15 mL sampling tubes (**do not leave any biomass behind**).
20. Centrifuge (5 min, 2500 x g, -19 °C).
21. Decant the supernatant into a waste vessel. **Important**: if you process samples for which you want to use internal standards for quantification of intracellular metabolites, internal standards are added at this step. See Basic Protocol 8.
22. Add 5 mL of 75% (v/v) ethanol (75 °C).
23. Resuspend the pellet by vortexing.
24. Close the sampling tubes using the marbles. *This prevents lids from popping off*.
25. Boil the samples 6 minutes by placing them in the 95 °C water bath.
26. Place the tubes on ice.
27. **Optional**: at this stage samples can be stored at -80 °C <u>up to 4 days</u>.
28. Evaporate the ethanol in the SpeedVac following manufacturer’s instructions. This typically takes approximately 12 hours. *It is recommended to execute the programme at room temperature. Do not heat to prevent breakdown of metabolites*.
29. After evaporation, resuspend the pellet in MilliQ. Adjust the MilliQ volume to reach 10 mg biomass per 100 μL of final volume. *The amount of biomass in the sample can be determined based on the dry weight measurement taken at the end of the fed-batch during the trial run. 10 mg biomass per 100 μL final volume was chosen based on previous research that found that this concentration should be sufficient to measure central carbon metabolism intermediates (Canelas et al., 2009; Mashego et al., 2004; Wasylenko & Stephanopoulos, 2015)*.
30. Centrifuge the resuspended cell extracts to remove cell debris (17000 x g, 5 min, 4 °C). Recover the supernatant that contains the cell extract and discard the pellets.
31. Centrifuge the recovered supernatant once more (17000 x g, 5 min, 4 °C) to ensure removal of all cell debris considering the high biomass concentration.
32. **Important**: when preparing internal standards, pool all resuspended cell extracts from all samples in a single flask to obtain a homogenous batch of ^13^C yeast extract.
33. Filter the (pooled) cell extract using 0.2 μm syringe filters.
34. Depending on the type of analysis to perform, proceed as follows:

a. For analysis of the ^12^C yeast extract from the trial run proceed to Basic Protocol 3 and 4.
b. For characterization of the ^13^C yeast extract continue to Basic Protocol 6 and 7.
c. For preparation of the internal standard only, divide the ^13^C yeast extract in screw-cap vials (1-2 mL of extract per vial) and store at -80 °C. **IMPORTANT**: if metabolites of interest are not present or insufficiently abundant in the ^13^C yeast extract, spike the internal standard with commercial ^13^C-labelled compounds at this stage, before dividing over vials.
d. For quantification of metabolites in a sample of interest using internal standard continue to Basic Protocol 8.

## Basic Protocol 3: Identification of intracellular metabolites using LC-MS

Metabolites that are polar and contain phosphate groups, like nucleotides, are best separated by LC using a porous graphitic carbon stationary phase (PGC). This protocol describes how to prepare samples to separate these compounds using PGC liquid chromatography and how to identify them using high resolution MS with metabolites involved in the NAD(P)^+^ biosynthesis in *S. cerevisiae* as example.

### Materials

Ammonium bicarbonate (Sigma Aldrich)

Optima LC/MS-grade H_2_O (Fisher Chemical), an aliquot of 50 mL is kept in the fridge

Optima LC/MS-grade acetonitrile (Fisher Chemical)

Mobile phase A: 50 mM ammonium bicarbonate in water (see reagents and solutions).

Mobile phase B: 100% Optima LC/MS-grade acetonitrile (Fisher Chemical)

Strong wash: 10% acetonitrile, 10 mM ammonium bicarbonate in water (see reagents and solutions).

Seal wash: 10% acetonitrile, 10 mM ammonium bicarbonate in water (see reagents and solutions).

Weak wash: 50 mM ammonium bicarbonate in water (see reagents and solutions).

Pierce™ LTQ ESI negative ion calibration solution (Thermo Fisher Scientific)

10 mM ammonium bicarbonate in 60% acetonitrile (see reagents and solutions).

Hypersep 96 Hypercarb 25/1ml Plate (Thermo Scientific)

2 ml Square Collection Plate, Part #186002482 (Waters)

Micro Centrifuge, speed 6000 rpm (Roth) or any other centrifuge for Eppendorf tubes

Collecting tubes, strips of 8 tubes (# SIMPT110-15, Avantor)

Acquity Ultra Performance Liquid Chromatograph (Waters)

Q Exactive™ Focus Hybrid Quadrupole-Orbitrap™ Mass Spectrometer (Thermo Scientific)

Chromatography column: Hypercarb™ Porous Graphitic Carbon HPLC Column, 100 x 1 mm, Thermo Scientific,

Part no. 35005-101030, particle size 5 μm

LC/MS vials: Polypropylene 12 x 32 mm Screw Neck Vial, with Cap and Preslit PTFE/Silicone Septum, 300 µL

Volume (Waters)

Glass 500 μL syringe for HPLC instruments (Thermo Scientific)

Analysis software (e.g. Xcalibur 4.1, Thermo Scientific)

12C commercial compounds of interest, highest purity

^12^C yeast cell extract as produced in Basic Protocol 1 and 2.

*CAUTION:* Electrical grounding of the porous graphitic carbon column is required to prevent column polarization caused by the electrospray voltage (Pabst & Altmann, 2008).

*CAUTION:* Retention time (RT) instability is often observed when using porous graphitic carbon as the chromatographic separation phase. Washing steps and regular backflushing could solve the problem (Bapiro et al., 2016). In this study, a mobile phase A, containing 50 mM ammonium bicarbonate, was used to limit the RT variability. Nevertheless, RT instability should be kept in mind and should be evaluated when performing the analyses. If necessary, inject the standards again separately or as a mixed solution prior to the analyses of the yeast cell extract.

*NOTE:* Whenever water is mentioned, it refers to Optima LC/MS-grade water.

### Protocol

#### Cleaning up the cell extracts using Solid Phase Extraction (SPE)

1. Wash and concentrate the cell extract obtained as described in step 34 of Basic Protocol 2, using a HyperSep PGC column plate as follows:

- Place a waste tray under the SPE columns.
- Condition step: Add 600 μL 10 mM ammonium bicarbonate in 60% acetonitrile to the SPE columns.
- Equilibration step: Add 2x 600 μL water to the SPE columns.
- Loading step: Pipette the samples onto the SPE columns.
- Washing step: Add 1000 μL water to the SPE columns.
- Remove the waste tray and place the sample collection tray with new collecting cups underneath the SPE columns.
- Elute the samples by adding 2 x 200 μL 10 mM ammonium bicarbonate in 60% acetonitrile to the SPE columns.

2. Transfer the samples from the collecting cups to 1.5 mL Eppendorf tubes.

3. Dry the collected samples using the SpeedVac concentrator on the “no temperature” setting. Leave the lid of the Eppendorf tubes open. This step usually takes three to four hours.

4. **Optional:** store the dried samples at -80 ᵒC until analysis.

5. Dissolve the samples in 20 μL of cold water.

6. Vortex the samples for a couple of minutes.

7. Gather all liquid at the bottom of the Eppendorf tubes by shortly centrifuging the samples (∼30 sec) using a table centrifuge.

*After centrifugation of the samples, you might notice a small black pellet in the tube. During elution from the PGC SPE columns, part of the graphite carbon might come into some samples. Avoid pipetting this black pellet into your MS vial, only take the clear supernatant*.

8. Transfer the supernatant to a MS vial. The samples can now be analysed by LC-MS.

#### Identifying metabolites using LC-MS

9. Prepare separate solutions of commercial compounds of interest dissolved in water, using the concentrations shown in Table 2.

10. Perform a mass calibration in negative mode using 500 μL Pierce™ LTQ ESI negative ion calibration solution.

11. Create a full MS method using the following settings:

- Method duration: 31 minutes
- Polarity: Negative
- Resolution: 35000
- #Scan ranges: 1
- Scan range: 75 to 1000 m/z
- Automatic Gain Control (AGC) target: 1.00E+06
- Maximum injection time (IT): 100 ms
- Microscans: 2
- Spectrum data type: Profile

12. Carefully purge the HPLC system with 50% mobile phase A (50 mM ammonium bicarbonate in optima LC/MS grade H_2_O) and 50% mobile phase B (Optima LC/MS grade acetonitrile) for 10 minutes and equilibrate the mounted and grounded PGC column overnight in the starting conditions.

13. Create a HPLC method using the following settings:

- Flow rate: 0.100 ml/min
- Set the gradient as described in Table 1
- Seal wash: 2.5 min

**Table 1.** Applied gradient for separation of metabolites by liquid chromatography. The mobile phase A consists of 50 mM ammonium bicarbonate in Optima LC/MS-grade H2O. The mobile phase B consists of Optima LC/MS-grade acetonitrile. Curve 6 means linear.

| Time (min) | %A | %B | Curve |
| --- | --- | --- | --- |
| Initial | 97.5 | 2.5 | Initial |
| 2.0 | 97.5 | 2.5 | 6 |
| 20.0 | 65 | 35 | 6 |
| 26.0 | 40 | 60 | 6 |
| 27.0 | 97.5 | 2.5 | 6 |
| 31.0 | 97.5 | 2.5 | 6 |

**Table 2.** Concentrations of commercial compounds of interest in the calibration solutions. Concentrations were rounded to whole number.

| Compound | Concentration (mM) | Compound | Concentration (mM) |
| --- | --- | --- | --- |
| PRPP | 13 | NR | 27 |
| ATP | 13 | NAM | 27 |
| ADP | 27 | NA | 20 |
| NADPH | 13 | Trp | 16 |
| NADP | 27 | QA | 1 |
| NADH | 27 | NMN | 27 |
| NAD | 27 | NaMN | 13 |
| NaAD | 27 |  |  |

PRPP: 5-phosphoribosyl-1-pyrophosphate; ATP: adenosine triphosphate; ADP: adenosine diphosphate; NAD(P)(H): nicotinamide adenine dinucleotide (phosphate); NaAD: nicotinic acid adenine dinucleotide; NR: nicotinamide riboside; NAM: nicotinamide; NA: nicotinic acid; Trp: tryptophan; QA: quinolic acid; NMN: nicotinamide mononucleotide; NaMN: nicotinic acid mononucleotide.

14. When creating the sample list, set the injection volume to 1 μL and run two blanks, consisting of Optima LC/MS grade H_2_O, in between each sample.

15. Place the solutions to analyse (cell extract or commercial compound) in the autosampler and start the run.

16. Analyse the mass spectra. This was performed in Xcalibur 4.1 (Thermo Scientific), but any other suitable software can also be used.

*This example focuses on the identification of NAD^+^-related metabolites, but many other metabolites can be identified if commercial compounds are available for calibration*.

*Extracted ion chromatograms were created of the metabolites of interest using a 10 ppm error range, after which the area of the peaks was integrated*.

***IMPORTANT***: This identification step determines which metabolites can be measured in the ^12^C yeast cell extract, and hence in the internal standard. Some metabolites might not be measurable due to several factors such as concentration below detection limit or chemical instability. For instance, nicotinic acid, although stable, is close to detection limit in yeast cell extracts. The yeast-based internal standard can therefore not be used for nicotinic acid quantification. This caveat can be mitigated by addition of commercial, ^13^C-labelled nicotinic acid to the custom-made internal standard.

17. Determine the retention time of each standard. The m/z of each unlabelled metabolite (negatively charged) is provided in Table 3.

**Table 3.** Selection of metabolites in NAD^+^ and ATP metabolism detectable by LC-MS with their corresponding masses for ^12^C and ^13^C variants.

| Compound | Monoisotopic mass | Chemical formula [M-H]<sup>-</sup> | $^{12}\text{C}$ $m/z$ | $^{13}\text{C}$ $m/z$ |
| --- | --- | --- | --- | --- |
| PRPP | 389.9518 | (C<sub>5</sub>H<sub>12</sub>O<sub>14</sub>P<sub>3</sub>)<sup>-</sup> | 388.9445 | 393.9613 |
| ATP | 506.9958 | (C<sub>10</sub>H<sub>15</sub>N<sub>5</sub>O<sub>13</sub>P<sub>3</sub>)<sup>-</sup> | 505.9885 | 516.0220 |
| ADP | 427.0294 | (C<sub>10</sub>H<sub>14</sub>N<sub>5</sub>O<sub>10</sub>P<sub>2</sub>)<sup>-</sup> | 426.0221 | 436.0557 |
| NADPH | 745.0911 | (C<sub>21</sub>H<sub>29</sub>N<sub>7</sub>O<sub>17</sub>P<sub>3</sub>)<sup>-</sup> | 744.0838 | 765.1543 |
| NADP | 744.0833 | (C<sub>21</sub>H<sub>27</sub>N<sub>7</sub>O<sub>17</sub>P<sub>3</sub>)<sup>-</sup> | 742.0682 | 763.1386 |
| NADH | 665.1248 | (C<sub>21</sub>H<sub>28</sub>N<sub>7</sub>O<sub>14</sub>P<sub>2</sub>)<sup>-</sup> | 664.1175 | 685.1880 |
| NAD | 664.1164 | (C<sub>21</sub>H<sub>26</sub>N<sub>7</sub>O<sub>14</sub>P<sub>2</sub>)<sup>-</sup> | 662.1019 | 683.1723 |
| **NaAD** | 665.1004 | (C<sub>21</sub>H<sub>25</sub>N<sub>6</sub>O<sub>15</sub>P<sub>2</sub>)<sup>-</sup> | 663.0859 | 684.1563 |
| **NR** | 255.0976 | (C<sub>11</sub>H<sub>13</sub>N<sub>2</sub>O<sub>5</sub>)<sup>-</sup> | 253.0830 | 264.1199 |
| **NAM** | 122.0480 | (C<sub>6</sub>H<sub>5</sub>N<sub>2</sub>O)<sup>-</sup> | 121.0407 | 127.0609 |
| **NA** | 123.0320 | (C<sub>6</sub>H<sub>4</sub>NO<sub>2</sub>)<sup>-</sup> | 122.0248 | 128.0449 |
| **Trp** | 204.0899 | (C<sub>11</sub>H<sub>11</sub>N<sub>2</sub>O<sub>2</sub>)<sup>-</sup> | 203.0826 | 214.1195 |
| **QA** | 167.0219 | (C<sub>7</sub>H<sub>4</sub>NO<sub>4</sub>)<sup>-</sup> | 166.0146 | 173.0381 |
| **NMN** | 334.0566 | (C<sub>11</sub>H<sub>14</sub>N<sub>2</sub>O<sub>8</sub>P)<sup>-</sup> | 333.0493 | 344.0862 |
| **NMN-Fragment** | 212.0086 | (C<sub>5</sub>H<sub>8</sub>O<sub>7</sub>P)<sup>-</sup> | 211.0008 | 216.0181 |
| **NaMN** | 335.0406 | (C<sub>11</sub>H<sub>13</sub>NO<sub>9</sub>P)<sup>-</sup> | 334.0333 | 345.0702 |

18. Identify metabolites of interest in the cell extract by comparing mass and retention time with the values obtained with the unlabelled commercial compounds.

*In our analysis, PRPP and QA commercial compounds could be detected, however, they did not retain well on the PGC column and eluted almost immediately. In the ^12^C yeast cell extract, PRPP and QA were below detection limit and could not be identified by LC-MS*.

## Basic Protocol 4: Identification of intracellular metabolites using GC-MS

Metabolites that can become volatile and are thermally stable are best separated by gas chromatography. Metabolites of central carbon metabolism (e.g. glycolysis, TCA cycle and the pentose phosphate pathway) and amino acids are such metabolites, they however require chemical modification to meet the requirements of GC-MS based separation and identification. Cell extracts are therefore freeze-dried and derivatized before GC-MS analysis. Glycolytic intermediates undergo oximation and silylation to form so-called TMS-MOX derivatives. Amino acids are silylated to obtain so-called TBDMS derivatives. Once the metabolites of interest are successfully identified in the ^12^C yeast cell extract, ^13^C-labelled yeast extract can be produced using Basic Protocol 5: Real run – Fed-Batch.

### Materials

Compounds of interest, p.a. or higher purity

Ultrapure water

Acetonitrile (HPLC grade)

Alkane standard, saturated C7-C30, 1000 μg/mL in hexane

Ethyl acetate (HPLC grade)

Hexane (HPLC grade)

Isooctane (HPLC grade)

Methoxyamine hydrochloride (MOX)

MSTFA; *N*-Trimethylsilyl-*N*-methyl trifluoroacetamide

MTBSTFA; N-methyl-N-(tert-butyldimethylsilyl)trifluoroacetamide with 1% tert-Butyldimethylchlorosilane

NaCl, 99% pure

Pyridine (HPLC grade)

Trimethylchlorosilane (TMCS) (BGB Analytik)

Pipettes

Clear glass deactivated, (silanized) 2 mL vial (Agilent)

Short thread screw caps (blue) with PTFE/silicone/PTFE septa (Agilent)

Short thread screw caps (transparent) with PTFE (Teflon) septa only (BGB Analytik) Snap top shell vial, 1 mL, 8 mm, clear (Agilent)

Vial inserts, 250 µL, pulled point glass (Agilent)

Freezer (-80 ⁰C)

Freeze dryer

Heating module (70 ⁰C)

Centrifuge

GC system: 7890 A (Agilent Technologies) or equivalent

MS: Triple Quad 7000 (Agilent Technologies) or equivalent

Zebron 50 ZB-capillary column (30 m x 250 µm internal diameter, 0.25 µm film) (Phenomenex)

Straight glass liner. Ultra Inert, single taper with glass wool (Agilent)

10 μL gastight syringe with Teflon-tipped plunger (Agilent)

***IMPORTANT***: For GC vials, silicone rubber should be avoided because of possible sample contamination and reactivity with the derivatization reagents.

### Protocol

#### Preparation of calibration solutions with unlabelled commercial compounds

1. Prepare stock solutions with commercial ^12^C compounds of interest in ultrapure water.

*During analysis, the concentration of commercial compounds should be in the same range as expected in the sample to analyse. A typical commercial calibration solution has concentrations between 0.25 μM and 100 μM when optimized for* S. cerevisiae*, however, depending on the microorganism of interest this concentration range might have to be adjusted. Use the certificate of analysis from chemical providers for preparation of stock solutions*.

2. From these stock solutions, prepare a calibration mix containing all commercial compounds (as example see Basic Protocol 7, Tables 10 and 11).

3. Prepare aliquots from this calibration mix and store at -80 °C.

4. For preparation of a calibration range with these commercial compounds see Basic Protocol 7.

#### Lyophilization of the cell extracts

5. Transfer 100 µL of cell extract or commercial compound to a silanized glass vial.

6. For amino acid analysis (and <u>not</u> for TCA and glycolysis metabolites) add 30 µL of 100 mg/mL NaCl in ultrapure water to each glass vial.

7. Close the vials with the caps containing white Teflon septa and puncture two holes in the septa with a needle.

8. Freeze for at least 20 min at –80 °C.

9. Place the frozen samples in a freeze dryer chamber.

*By freezing samples in a 45° angle more surface area is created for lyophilization*.

10. Start the vacuum pump.

11. Place the samples in the freeze dryer overnight.

12. After freeze-drying recap the vials with blue caps with PTFE/silicone/PTFE septa.

13. Store samples at –80 °C until derivatization.

#### Derivatization of metabolites

*CAUTION:* Work in the fume hood and wear nitrile gloves for the derivatization.

**<u>For derivatization of central carbon metabolites</u>** (glycolysis, pentose phosphate pathway, and TCA metabolites)

14. Switch on the heating block (70 °C).

15. Prepare a MSTFA silylation reagent mix: Mix 1 mL of MSTFA with 50 μL TMCS. Use p.a. or higher purity.

*Derivatizing reagents should be stored in a desiccator in a fridge at 4 °C*.

16. Prepare a MOX solution of 20 mg/mL methoxyamine in anhydrous pyridine. Vortex and preferably heat for 1 min in the heating block to let MOX completely dissolve (prepare freshly every day).

17. Add 50 μL of the MOX solution to each GC vial.

18. Incubate cell extracts or commercial compounds for 50 min at 70 °C in the heating block.

19. Remove vials from the heating block and let cool down to room temperature.

20. Add 80 μL of the MSTFA silylation reagent mix to each vial.

21. Incubate the vials for 50 min at 70°C in the heating block.

22. Remove vials from the heating block and let them cool down to room temperature.

23. Transfer samples from vials to centrifuge glass tubes (1 mL shell vials).

24. Centrifuge (1 min, 10,000 x g, 4°C).

25. Transfer 90 μL of supernatant from the centrifuge tube into a glass insert and place it back into the original GC vial (remove air bubbles if needed).

#### For derivatization of amino acids

26. Switch on the heating block (70 °C).

27. Add 75 μL of acetonitrile and 75 μL MTBSTFA to each GC vial.

28. Incubate samples for 60 min at 70 °C in the heating block.

29. Remove vials from the heating block and let them cool down to room temperature.

30. Transfer samples into the centrifuge glass tubes (1 mL shell vials).

31. Centrifuge (2 min, 10,000 x g, 4°C).

32. Transfer 90 μL of supernatant from the centrifuge tube into a glass insert and place it back into the original GC vial (remove air bubbles if needed).

*After these derivatization steps, the samples are ready for analysis by GC-MS. The derivatized samples can be kept in the fridge for up to two weeks*.

#### Analysing the derivatized metabolites from central carbon metabolism by GC-MS (according to Niedenführ et al. (2016) and Seifar et al. (2018))

33. Set the carrier gas flow (helium 99.9990%) to 1 mL/min.

34. Program the GC oven temperature as follows: keep constant for 1 min at 70 °C followed by a ramp of 10 °C per min up to 220 °C.

35. Set the temperature of the transfer line to 250 °C.

36. Set the injection volume to 1 μL.

37. If using a multimode inlet (MMI) (Agilent), use the following settings: split-less mode for metabolites with low concentration and in split mode 1:5 for metabolites with high concentrations.

38. Set the MMI temperature: 70 °C (with a split-less time of 1 min if it is needed), after injection temperature is increased to 220 °C with a ramp of 720 °C/min and held for 5 min. Then the temperature is increased to 300 °C to clean the injector.

39. At the end of the run, backflush the column with five column volumes at 220 °C.

*For multiple injections from the same sample vial, renew the cap after each two injections if it is possible*.

*The liner in the GC might need to be replaced based on the sample type after different numbers of injections. After about 100 injections, cut 10 cm of the column (injection side) and at the same time replace the liner and inlet septum (gas flow needs to be modified accordingly using retention time locking (RTL), to have reproducible retention times*.

*Run the alkane mix (20 µg/ml saturated alkane C7-C30 in hexane) for RTL and system check about every ten samples*.

40. Set the MS transfer line to 250 °C.

41. Keep the MS source at 230 °C.

42. Set the helium gas flow of the collision cell to 2.25 mL/min.

43. Set the N2 collision gas flow to 1.5 mL/min.

44. Use electron ionization operated at 70 eV.

45. The MS is operated in SRM mode.

46. Mass resolution is set to 0.7 mass unit for both quadrupoles.

47. Run GC-MS analysis

48. Mass Hunter analysis software (version 12.0/12.1; Agilent) can be used for data processing.

49. Integrate signals and qualify compounds using the commercial standard mix as a reference.

50. Identify all the compounds of interest in the unlabelled yeast extract sample based on the properties determined using the commercial compounds mix (Table 4).

**Table 4.** The m/z values, number of C-atoms, and collision energy of some central carbon metabolites of interest.

| Compound | Precursor m/z | Original number of C-atoms in precursor | Product m/z | Original number of C-atoms in product | Collision Energy |
| --- | --- | --- | --- | --- | --- |
| **Fum** | 245.0 | 4 | 146.9 | 0 | 10 |
| **Succ** | 247.0 | 4 | 147.1 | 0 | 5 |
| **Mal** | 335.0 | 4 | 146.9 | 0 | 15 |
| **aKG** | 304.0 | 5 | 146.9 | 0 | 5 |
| **G3P** | 445.1 | 3 | 299.2 | 0 | 15 |
| **GAP** | 400.1 | 3 | 299.0 | 0 | 10 |
| **Glc** | 319.1 | 4 | 129.1 | 3 | 10 |
| **Cit** | 465.1 | 6 | 146.9 | 0 | 35 |
| **iCit** | 465.1 | 6 | 146.9 | 0 | 35 |
| **2PG** | 459.1 | 3 | 299.1 | 0 | 15 |
| **DHAP** | 400.1 | 3 | 299.1 | 0 | 10 |
| **3PG** | 459.1 | 3 | 299.1 | 0 | 15 |
| **E4P** | 357.1 | 2 | 341.1 | 2 | 15 |
| **Rib5P** | 459.1 | 3 | 315.2 | 0 | 10 |
| **Ribu5P** | 357.1 | 2 | 147.1 | 0 | 35 |
| **Xyl5P** | 357.0 | 2 | 147.1 | 0 | 30 |
| **M6P** | 471.0 | 4 | 387.0 | 0 | 10 |
| **F6P** | 459.1 | 3 | 315.2 | 0 | 10 |
| **G6P** | 471.0 | 4 | 387.0 | 0 | 10 |
| **6PG** | 333.1 | 4 | 73.1 | 3 | 10 |
| **S7P** | 471.1 | 4 | 387.1 | 0 | 10 |
| **Tre** | 361.1 | 6 | 169.1 | 5 | 15 |
| **FBP** | 459.2 | 3 | 315.0 | 0 | 10 |
| **T6P** | 513.1 | 6 | 315.1 | 0 | 10 |

*Fum: fumarate; Succ: succinate; Mal: malate; aKG: α-ketoglutarate; G3P: glycerol-3-phosphate; GAP: glyceraldehyde-3-phosphate; Glc: glucose; Cit: citrate; iCit: iso-citrate; 2PG: 2-phosphoglycerate; 3PG: 3-phosphoglycerate; DHAP: dihydroxyacetone phosphate; E4P: erythrose-4-phosphate; Rib5P: ribose-5-phosphate; Ribu5P: ribulose-5-phosphate; Xyl5P: xylulose-5-phosphate; M6P: mannose-6-phosphate; F6P: fructose-6-phosphate; G6P: glucose-6-phosphate; 6PG: 6-phosphogluconate; S7P: sedoheptulose-7-phosphate; Tre: trehalose; FBP: fructose-1,6-bisphosphate; T6P: trehalose-6-phosphate.*

#### Analysing derivatized amino acids by GC-MS (according to De Jonge et al. (2011))

51. Set the carrier gas flow (helium 99.9990%) to 1 mL/min.

52. Program the GC oven temperature as follows: keep constant for 1 min at 100 °C followed by a ramp of 10 °C per min up to 320 °C.

53. Set the temperature of the transfer line to 250 °C.

54. Set the injection volume to 1 μL.

55. If using a multimode inlet (MMI) (Agilent), use the following settings: split-less mode for metabolites with low concentration and in split mode 1:5 for metabolites with high concentrations.

56. Set the MMI temperature: 100 °C (with a split-less time of 1 min if it is needed), after injection temperature is increased to 220 °C with a ramp of 720 °C/min and held for 5 min. Then the temperature is increased to 300 °C to clean the injector.

57. At the end of the run, backflush the column with five column volumes at 320 °C.

*The same comments for step 39 in this protocol also apply to step 57*.

58. Set the MS transfer line to 250 °C.

59. Keep the MS source at 230 °C and the quadrupole at 150°C.

60. Use electron ionization operated at 70 eV.

61. The MS is operated in SIM mode.

62. Mass resolution is set to 1 mass unit throughout the mass range 1-1050 amu.

63. Run GC-MS analysis

64. Mass Hunter analysis software (version 12.0/12.1; Agilent) can be used for data processing.

65. Identify all the amino acids of interest in the unlabelled yeast cell extract based on the properties defined for the prepared commercial compounds mix (Table 5).

**Table 5.**
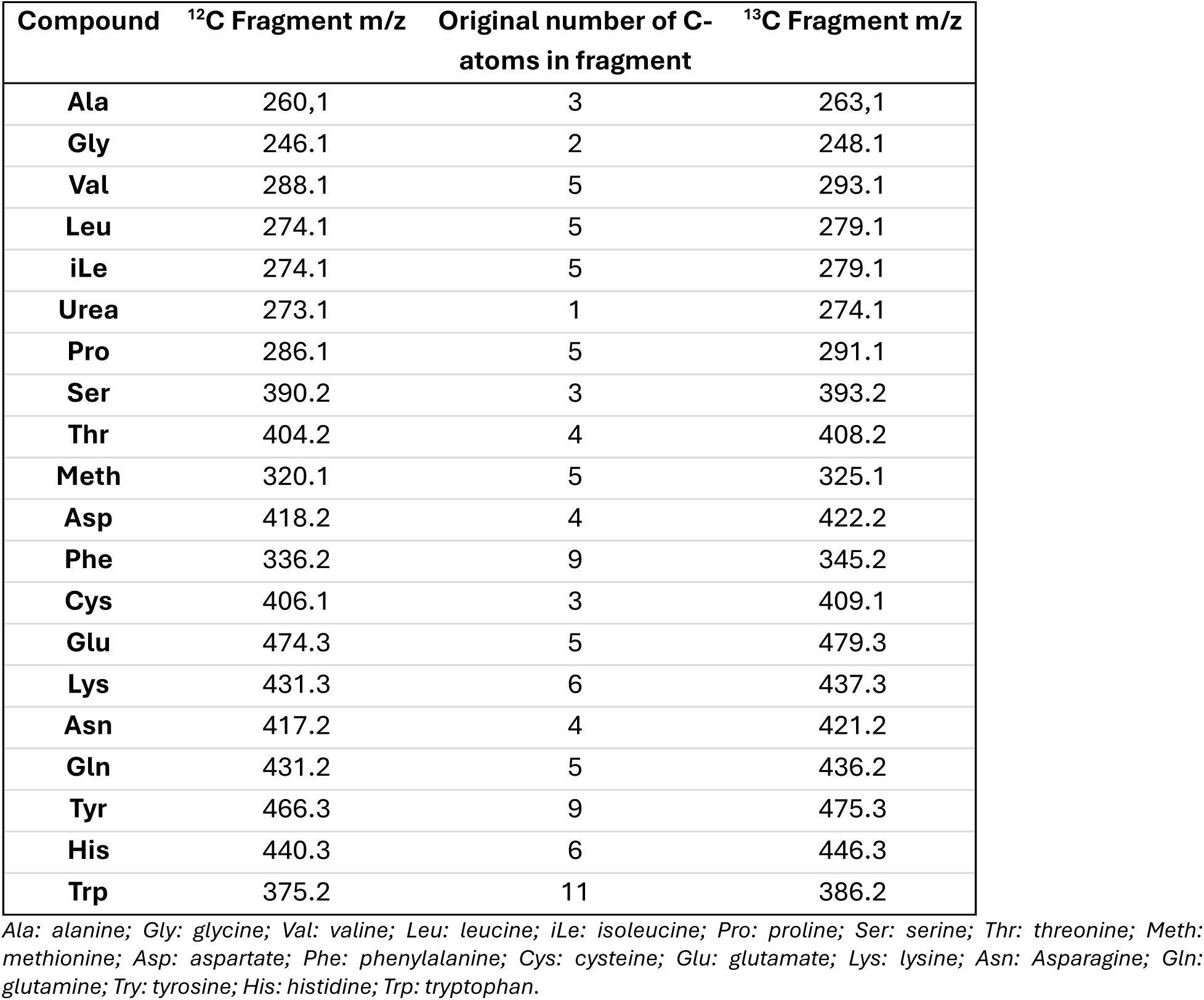
m/z values, number of C-atoms of amino acids.

| **Compound** | **<sup>12</sup>C Fragment m/z** | **Original number of C-atoms in fragment** | **<sup>13</sup>C Fragment m/z** |
| --- | --- | --- | --- |
| **Ala** | 260,1 | 3 | 263,1 |
| **Gly** | 246.1 | 2 | 248.1 |
| **Val** | 288.1 | 5 | 293.1 |
| **Leu** | 274.1 | 5 | 279.1 |
| **iLe** | 274.1 | 5 | 279.1 |
| **Urea** | 273.1 | 1 | 274.1 |
| **Pro** | 286.1 | 5 | 291.1 |
| **Ser** | 390.2 | 3 | 393.2 |
| **Thr** | 404.2 | 4 | 408.2 |
| **Meth** | 320.1 | 5 | 325.1 |
| **Asp** | 418.2 | 4 | 422.2 |
| **Phe** | 336.2 | 9 | 345.2 |
| **Cys** | 406.1 | 3 | 409.1 |
| **Glu** | 474.3 | 5 | 479.3 |
| **Lys** | 431.3 | 6 | 437.3 |
| **Asn** | 417.2 | 4 | 421.2 |
| **Gln** | 431.2 | 5 | 436.2 |
| **Tyr** | 466.3 | 9 | 475.3 |
| **His** | 440.3 | 6 | 446.3 |
| **Trp** | 375.2 | 11 | 386.2 |

*Ala: alanine; Gly: glycine; Val: valine; Leu: leucine; iLe: isoleucine; Pro: proline; Ser: serine; Thr: threonine; Meth: methionine; Asp: aspartate; Phe: phenylalanine; Cys: cysteine; Glu: glutamate; Lys: lysine; Asn: Asparagine; Gln: glutamine; Try: tyrosine; His: histidine; Trp: tryptophan.*

## Basic Protocol 5: Real run – Fed-batch

When the fed-batch fermentation reached (near) maximum biomass yield on glucose, meaning mainly respiratory metabolism for *S. cerevisiae*, and the ^12^C yeast extract contained (most of) the metabolites of interest, you are ready for the ‘real run’. To obtain a fully ^13^C-labelled metabolite extract, a few deviations of Basic Protocol 1 are required at the following steps:

Step 2. For the preparation of the batch, add the ^13^C glucose solution and be aware that the ^13^C glucose solution requires a slightly different amount of glucose (see Reagents and Solutions).

Step 5. The sampling at regular time intervals should be skipped. The trial run will inform the real run, and no further samples will be taken during the fermentation to prevent loss of valuable ^13^C yeast.

Step 6. As described in step 6, monitor the fermentation closely via gas analysis. It is essential to use a gas analyser that can detect ^13^C-CO_2_ as previously described, however, even those gas analysers will be less sensitive towards ^13^C-CO_2_ and as a result the signal will be lower compared to the trial run.

Step 7. The fed-batch feeding profile as calculated during the trial run will be used for the real run.

Step 9. During the fed-batch, step 9 will be skipped, and no samples will be taken during the fed-batch phase. If the RQ during the fed-batch phase is too high according to CO_2_ and DO data, the fed-batch profile should not be adjusted. During the real run, the data obtained is insufficient and less accurate to support an informed decision on adjusting the feed rate; therefore, the feeding profile determined during the trial run will be used.

For the sampling, no adjustments to Basic Protocol 1 are required. The ratio of methanol to broth and the timing of the sampling are kept the same. In Supplementary File 1, spreadsheet ‘FEDBATCH_SAMP’ nine sampling points are suggested, however, the number of sampling points can be adjusted based on lab capacity. These nine sampling timepoints were based on the lab capacity of one centrifuge, that can centrifuge six 50 mL Greiner tubes at a time, and -40 °C freezer capacity for six 50 mL Greiner tubes. In the case of this lab equipment, the leftover six tubes that cannot be immediately centrifuged should be kept in the -40 °C freezer. The most important during the sampling and subsequent sampling processing is to keep the samples cold, i.e. quenched, at all times.

Finally, sample processing as described in Basic Protocol 2 can be followed without any deviations.

## Basic Protocol 6: LC-MS characterization of the internal standard

Basic Protocol 6 describes the assessment of ^12^C contamination of the internal standard produced in Basic Protocol 5 using LC-MS, the identification of the metabolites of interest, and the quantification of these metabolites in the internal standard using unlabelled commercial standards.

### Materials

Same as Basic Protocol 3

### Protocol

#### Evaluation of potential contamination of the metabolites by ^12^C in the internal standard

While the culture medium has been optimized to minimize the occurrence of ^12^C in the internal standards, the purity of the ^13^C-labelled yeast extract needs to be evaluated. The number of ^12^C and ^13^C in a metabolite will affect its mass, which can be detected by mass spectrometry. The internal standard is therefore analysed by LC-MS as described in Basic Protocol 3.

Since the mass difference between a ^13^C and a ^12^C atom equals 1.003354, the mass of the isotope variants can be calculated using equation 3:

Isotope mass= monoisotopic mass+N∗1.003354 (Equation 3, N represents the number of carbon atoms)

For example, the ^12^C monoisotopic mass of ATP equals 506.99575 and ATP contains ten carbon atoms. Therefore, the mass of fully ^13^C-labelled ATP becomes: 506.99575+10∗1.003354=517.02929

The mass of ATP containing nine ^13^C atoms and one ^12^C atom becomes:

506.99575+9∗1.003354= 516.025936

Since, the MS analyses are performed in negative mode, the mass of a proton needs to be subtracted to get the final m/z value which can be used to create the extracted ion chromatograms, see Table 3.

1. Analyse the ^13^C yeast extract by following the instructions of Basic Protocol 3 steps 1 to 16.
2. Create for all isotope variants of each metabolite an extracted ion chromatogram with a 10 ppm error marge.
3. Integrate the peaks in Xcalibur or any other suitable software.
4. Calculate the percentage of isotope variants compared to the fully ^13^C-labelled variant using equation 4.

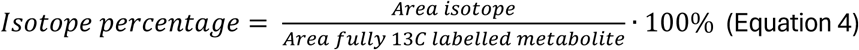

While full ^13^C-labelling of all carbon atoms in all metabolites is difficult to reach and is not required for the internal standards, the absence of ^12^C monoisotopes is a prerequisite to avoid interference between internal standard and unlabelled samples to analyse. For instance, the internal standard produced with this protocol contained metabolites with one ^12^C atom (5-28% of all metabolites), and to a lesser extent two (0-7%) or three (0-2%) ^12^C atoms. At least half of the carbon atoms in each analysed metabolite consisted of ^13^C. However, ^12^C monoisotopes were absent from the internal standard. The underlying calculations can be found in Supplementary File 4 ‘Check_for_12C_Contamination’.

#### Quantification of metabolite concentrations in the internal standard using commercial ^12^C compounds

5. Prepare calibration mixtures with ^12^C commercial compounds in duplicate according to Table 6.

**Table 6.** Concentration (μM) of the commercial compounds of interest in the six calibration mixes (S1 to S6).

|  | NaAD | NR | NMN | NAM | NAD | NADP | NADH | ADP | NA | NADPH | NaMN | ATP | Trp |
| --- | --- | --- | --- | --- | --- | --- | --- | --- | --- | --- | --- | --- | --- |
| **S1** | 0.1 | 1 | 5 | 5 | 500 | 50 | 50 | 250 | 25 | 10 | 1 | 500 | 50 |
| **S2** | 0.5 | 5 | 10 | 50 | 1000 | 100 | 100 | 500 | 50 | 25 | 5 | 1000 | 75 |
| **S3** | 1 | 10 | 25 | 100 | 2000 | 200 | 200 | 1000 | 100 | 50 | 10 | 2000 | 100 |
| **S4** | 2 | 20 | 50 | 250 | 3000 | 300 | 300 | 2500 | 150 | 75 | 15 | 3000 | 150 |
| **S5** | 3.5 | 35 | 75 | 400 | 4000 | 400 | 400 | 4000 | 200 | 120 | 20 | 4000 | 200 |
| **S6** | 5 | 50 | 100 | 500 | 5000 | 500 | 500 | 5000 | 250 | 150 | 25 | 5000 | 250 |

*Intracellular metabolite concentrations typically range from micro to millimolars. The chosen range of concentrations of commercial compounds for quantification of metabolites in the internal standard are therefore designed to span this range. If the concentration of metabolite in the internal standard deviates too much from this range, a new calibration should be performed for the concerned metabolites, with adjusted concentrations of commercial compounds in the higher or lower range*.

6. Spike each calibration mixture with 100 μL ^13^C-labelled yeast extract.

*The volume of spiked 13C-labelled yeast extract was optimized to meet the detection limit of the mass spectrometer. While the volume of internal standard used should be minimized to reduce cost, ^13^C-labelled metabolites should be clearly detectable by the mass spectrometer*

7. Analyse the samples as instructed in Basic Protocol 3 steps 1 to 16.

8. Create extracted ion chromatograms of the metabolites of interest using a +/- 10 ppm error marge.

9. Integrate the area of the ^12^C metabolites and of the ^13^C-labelled metabolites.

*For NMN, a strong signal for its in-source fragment was observed (m/z 211.0008 for the fully ^12^C-labelled variant and m/z 216.0181 for the fully ^13^C-labelled variant). Because this fragment yielded a stronger signal than the intact compound, it was selected for creating the calibration curve*.

10. Calculate the ^12^C/^13^C area ratio for each metabolite of interest and plot it against the known concentration of the ^12^C commercial standards (see example in Figure 6).

**Figure 6.**
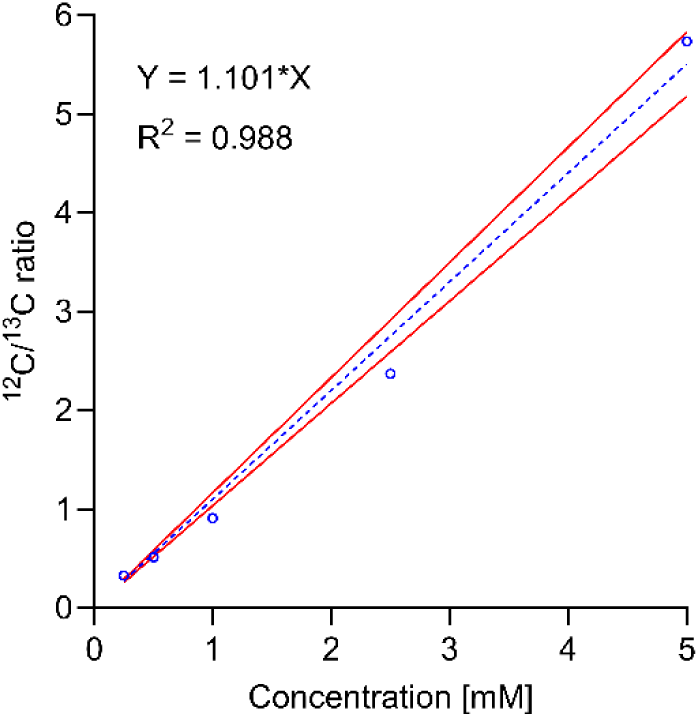
LC-MS-based quantification of ADP concentration in custom-made internal standard. The internal standard is mixed at fixed volume ratio with solutions of unlabelled commercial standards covering a broad concentration range (Table 7). The ratio of ^12^C over ^13^C high resolution mass spectrometry signals measured in these calibration mixtures is plotted against the known ADP concentration in the unlabelled commercial standards. The red lines represent the 95% confidence interval. This figure was generated using GraphPad Prism version 11.0.2.

**Table 7.** Linearity of the calibration curves over a specified concentration range.

| Compound | Concentration range ($\mu\text{M}$) | Linearity ($R^2$) |
| --- | --- | --- |
| ATP | 500-5000 | 0.956 |
| ADP | 250-5000 | 0.988 |
| NADPH | 10-150 | 0.956 |
| NADP | 50-500 | 0.968 |
| NADH | 50-500 | 0.916 |
| NAD | 500-5000 | 0.974 |
| NaAD | 0.1-5 | 0.987 |
| NR | 5-50 | 0.986 |
| NAM | 5-500 | 0.981 |
| NA | 25-250 | 0.989 |
| Trp | 50-250 | 0.965 |
| NMN | 5-100 | 0.988 |
| NaMN | 1-25 | 0.970 |

11. The linearity of this correlation for each metabolite can be calculated (Table 7).

12. Identify the concentration corresponding to a ^12^C/^13^C ratio of 1, which represents the metabolite concentration in the ^13^C yeast extract, see Table 8.

**Table 8.** Quantification of metabolite concentration in internal standards using calibration with commercial compounds. A 95% confidence interval is used.

| **Compound** | **Concentration (μM)** | **95% confidence interval range** | **RT (min)\*** |
| --- | --- | --- | --- |
| **ATP** | 2279.3 | 1986 – 2571 | 10.0 +/- 0.6 |
| **ADP** | 908.0 | 726 – 1076 | 9.9 +/- 0.4 |
| **NADPH** | 24.2 | 11 – 35 | 12.5 +/- 0.1 |
| **NADP** | 135.1 | 105 – 161 | 13.7 +/- 0.2 |
| **NADH** | 52.3 | 0 – 108 | 15.3 +/- 0.1 |
| **NAD** | 1151.5 | 865 – 1396 | 16.5 +/- 0.2 |
| **NaAD** | 0.4 | 0 – 1 | 13.2 +/- 0.2 |
| **NR** | 6.5 | 4 – 9 | 14.8 +/- 0.2 |
| **NAM** | 110.5 | 91 – 129 | 9.9 +/- 0.4 |
| **NA** | 123.5 | 117 – 130 | 5.4 +/- 0.7 |
| **Trp** | 78.4 | 63 – 91 | 15.0 +/- 0.2 |
| **NMN** | 11.6 | 8 – 15 | 7.5 +/- 0.1 |
| **NaMN** | 8.4 | 7 – 10 | 6.0 +/- 0.6 |

\*The reported retention time (RT) represents the average of the identified retention times for the fully <sup>13</sup>C-labelled metabolites across all calibration samples. The +/- value indicates the observed RT range.

*If a ratio of 1 lies outside the calibration curve, adjust the concentration range of the commercial ^12^C compounds (Table 6) and generate new calibration curves*.

*The linearity of most of the compounds identified in the internal standard is acceptable for metabolomics with a R^2^ of > 0.98 (Table 7). The linearity of NADH is rather low which might be due to the difficulty of measuring redox cofactors NAD(P)(H). As a result, we cannot confidently assign the NADH concentration to the internal standard (Table 8), however, the internal standard can still be used to show differences in NADH concentrations between conditions*.

## Basic Protocol 7: GC-MS characterization of the internal standard

Similar to Basic Protocol 6 using LC-MS, this protocol describes how to quantify the glycolytic intermediates and amino acid concentrations in the internal standard by GC-MS. Calibration curves with commercial ^12^C standards are used to quantify the concentration of each compound of interest in the internal standard. The lyophilization, derivatization, and GC-MS analysis are the same as for Basic Protocol 4. The deviation from Basic Protocol 4 is in the sample preparation step, and the final quantification of the internal standard. When Basic Protocol 7 is completed successfully, the internal standard is quantified and ready to be used as internal standard for future metabolomics studies on NAD(P) metabolites, see Basic Protocol 8.

### Materials

Same as Basic Protocol 4

### Protocol

1. Prepare stock solutions for each commercial compound in ultrapure water as instructed in step 1 of Basic Protocol 4).

2. From these stock solutions, prepare a calibration mix containing all targeted metabolites in known concentrations (Table 10) (Table 11).

*The concentrations in the calibration mix are the same as shown for CAL8 in Table 11*.

3. Prepare aliquots from this calibration mix and store at -80 °C.

4. Prepare calibration samples 1 to 10 (CAL1 – CAL10) following the pipetting scheme (Table 9). The range of concentrations of the ^12^C compounds is displayed in Table 10 and Table 11.

**Table 9.**
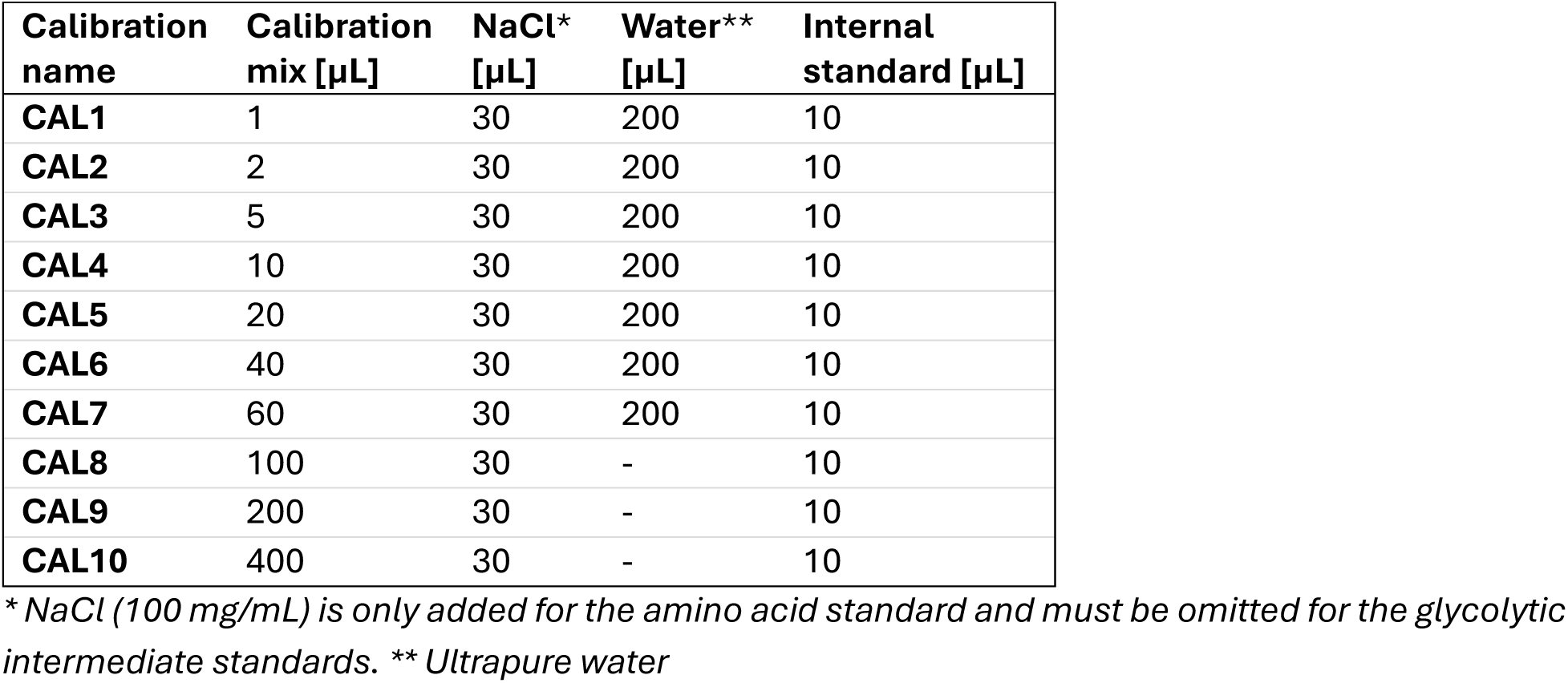
Pipetting scheme to obtain calibration mixes with a broad range of metabolite concentrations.

**Table 10.**
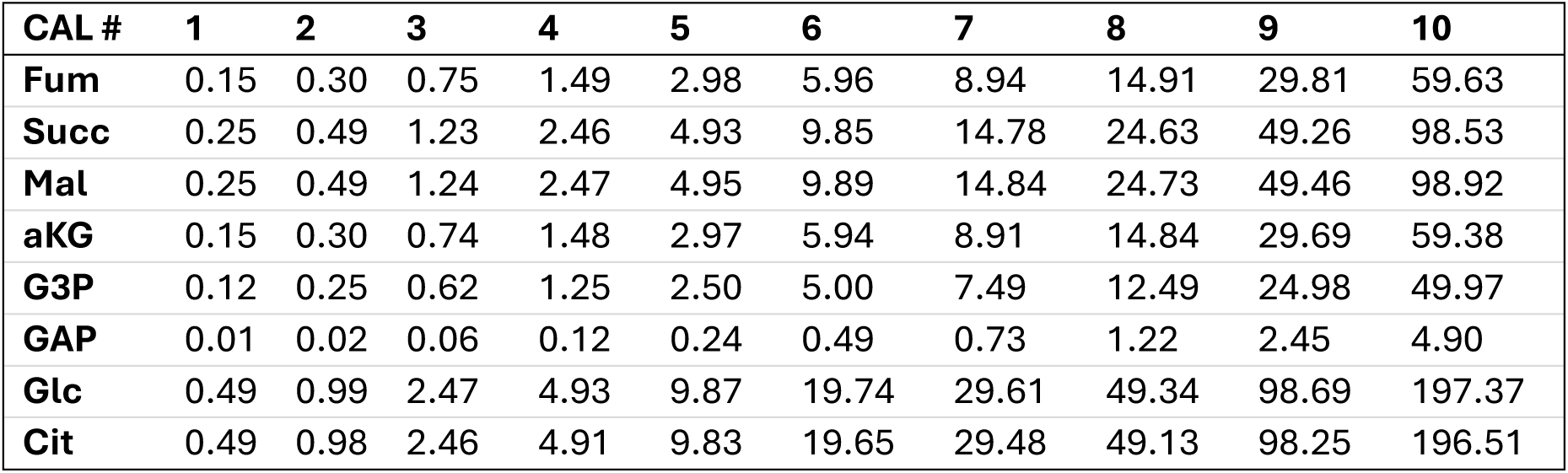
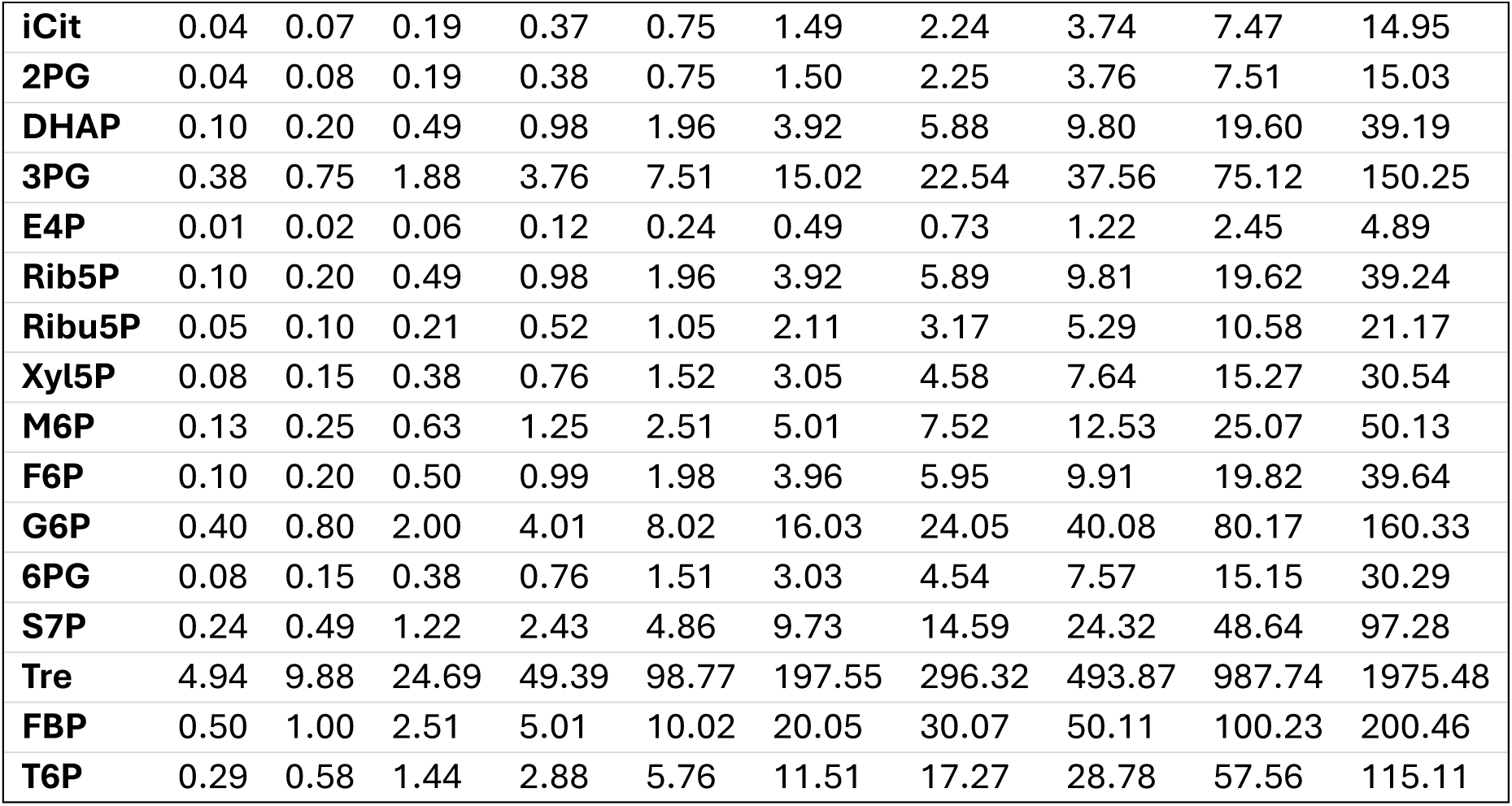
Concentrations of the ^12^C commercial compounds in the glycolytic intermediate calibration mixture. Concentrations are given in µM. CAL 8 corresponds to the concentration for the aliquoted calibration mixture.

**Table 11.** Concentrations of the ^12^C commercial compounds in the amino acid calibration mixture. Concentrations are given in µM. CAL 8 corresponds to the concentration for the aliquoted calibration mixture.

| **CAL #** | **1** | **2** | **3** | **4** | **5** | **6** | **7** | **8** | **9** | **10** |
| --- | --- | --- | --- | --- | --- | --- | --- | --- | --- | --- |
| **Ala** | 2.19 | 4.38 | 10.94 | 21.88 | 43.75 | 87.51 | 131.26 | 218.77 | 437.53 | 875.07 |
| **Gly** | 0.28 | 0.57 | 1.42 | 2.84 | 5.68 | 11.36 | 17.04 | 28.40 | 56.80 | 113.60 |
| **Val** | 0.51 | 1.01 | 2.53 | 5.06 | 10.12 | 20.25 | 30.37 | 50.62 | 101.24 | 202.48 |
| **Leu** | 0.06 | 0.12 | 0.30 | 0.60 | 1.20 | 2.40 | 3.60 | 6.00 | 12.00 | 24.00 |
| **Ile** | 0.06 | 0.12 | 0.31 | 0.61 | 1.22 | 2.44 | 3.66 | 6.11 | 12.21 | 24.43 |
| **Pro** | 0.18 | 0.37 | 0.91 | 1.83 | 3.65 | 7.30 | 10.95 | 18.25 | 36.50 | 73.01 |
| **Ser** | 0.51 | 1.02 | 2.54 | 5.09 | 10.17 | 20.35 | 30.52 | 50.86 | 101.73 | 203.45 |
| **Thr** | 0.51 | 1.01 | 2.54 | 5.07 | 10.15 | 20.29 | 30.44 | 50.73 | 101.46 | 202.91 |
| **Met** | 0.06 | 0.12 | 0.29 | 0.59 | 1.17 | 2.34 | 3.51 | 5.86 | 11.71 | 23.42 |
| **Asp** | 1.74 | 3.48 | 8.69 | 17.39 | 34.78 | 69.56 | 104.33 | 173.89 | 347.78 | 695.57 |
| **Phe** | 0.04 | 0.08 | 0.20 | 0.40 | 0.80 | 1.61 | 2.41 | 4.02 | 8.04 | 16.08 |
| **Cys** | 0.06 | 0.12 | 0.30 | 0.60 | 1.19 | 2.38 | 3.57 | 5.95 | 11.91 | 23.81 |
| **Glu** | 3.97 | 7.94 | 19.85 | 39.70 | 79.39 | 158.79 | 238.18 | 396.97 | 793.94 | 1587.88 |
| **Lys** | 0.24 | 0.48 | 1.19 | 2.38 | 4.76 | 9.52 | 14.28 | 23.81 | 47.62 | 95.23 |
| **Asn** | 0.25 | 0.49 | 1.23 | 2.46 | 4.91 | 9.83 | 14.74 | 24.57 | 49.14 | 98.28 |
| **Gln** | 2.03 | 4.06 | 10.14 | 20.28 | 40.56 | 81.12 | 121.68 | 202.79 | 405.59 | 811.18 |
| **Tyr** | 0.06 | 0.12 | 0.30 | 0.59 | 1.18 | 2.37 | 3.55 | 5.92 | 11.85 | 23.69 |
| **His** | 0.24 | 0.48 | 1.20 | 2.40 | 4.81 | 9.62 | 14.42 | 24.04 | 48.08 | 96.16 |
| **Trp** | 0.05 | 0.10 | 0.25 | 0.50 | 0.99 | 1.99 | 2.98 | 4.97 | 9.94 | 19.89 |

*Calibration samples were prepared by adjusting the added volume of calibration mix according to the target concentration (Table 9). Lower volumes were used for low concentration calibration samples (CAL 1-7), while higher volumes were used for high concentration calibration samples (CAL 9-10). The use of larger volumes is compatible with the subsequent lyophilization step, which removes excess solvent and standardizes final sample concentrations*.

*For calibration samples with an initial volume below 100 µL (corresponding to low concentrations of commercial compounds, Table 9), 200 µL of water was added prior to freezing to prevent defrosting during the freeze drying*.

5. Lyophilize samples CAL1-CAL10 (see Basic Protocol 4).

6. Derivatize samples CAL1-CAL10 (see Basic Protocol 4).

7. Analyse samples CAL1-CAL10 (see Basic Protocol 4).

8. Mass Hunter analysis software (version 12.0/12.1; Agilent) can be used for data processing.

9. Integrate signals and calculate concentrations based on the calibration line as established according to Table 9.

10. Calibration graphs are constructed based on peak area ratios of natural ^12^C peak area of targeted metabolites to ^13^C labelled peak area of internal standards versus concentration of the commercial natural isotope ^12^C compound in the calibration mixes. Use 1/y^2^ weighting for construction of calibration curves. The calibration curve for glucose-6-phosphate is shown in Figure 7 as example.

**Figure 7.**
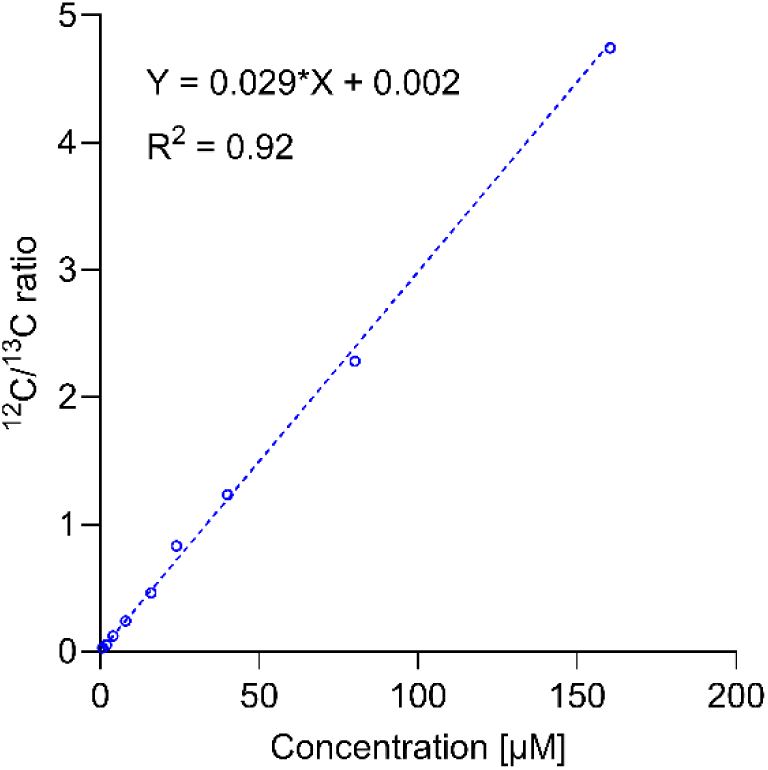
GC-MS-based quantification of glucose-6-phosphate concentration in the home-made internal standard. The internal standard is mixed at fixed volume ratio with solutions of unlabelled commercial standards covering a broad concentration range (Table 10). The ratio of ^12^C over ^13^C high resolution mass spectrometry signals measured in these calibration mixtures is plotted against the known glucose-6-phosphate concentration in the unlabelled commercial standards. Measurements were performed in analytical duplicates. The calibration curve was generated using non-linear regression with 1/y² weighting. This figure was generated using GraphPad Prism version 11.0.2.

11. The concentration of the compound in the ^13^C yeast extract is calculated from the calibration curves using equation 5.

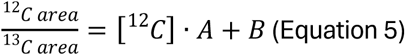

A: slope of the calibration curve

B: intercept of the calibration curve

12C area: peak area of unlabelled (^12^C) metabolite in the calibration mix

13C area: peal area of labelled (^13^C) metabolite in the calibration mix

[^12^C]: concentration of the ^12^C metabolite

Slope A of the calibration curve is obtained by linear regression giving equation 6.

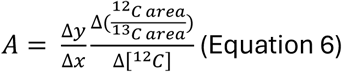

When working with an internal standard, the response factor of both the compound of interest and the internal standard is needed for the correction. However, as the ^12^C and ^13^C variants of a compound have the same chemical composition, their response factor (Rf) is the same. In this case Rf = 1.

The intercept B of the calibration curve reflects contributions from residual ^12^C metabolites or other matrix components present in the internal standard. Since we match the internal standard concentrations in both the calibration mixes and the samples, the relative contribution of ^12^C contamination and matrix effects is expected to be equivalent. Consequently, the intercept was assumed consistent and omitted to simplify the calculations.

The concentration of the ^13^C compounds in the internal standard were obtained using equation 5, 6, and 7.

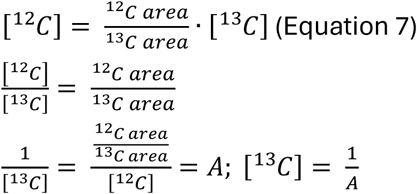

See Table 12 for concentrations of glycolytic intermediates and Table 13 for concentrations of amino acids in the internal standard. The concentrations are given for an amount of 100 µL internal standard.

**Table 12.** Quantification of glycolytic metabolite concentrations in the internal standard using calibration with commercial compounds. The reported concentration has been corrected for the addition of 10 µL internal standard to the calibration sample. All samples were measured in duplicate.

| Compound | Concentration [µM] | Slope | Intercept | Linearity (R<sup>2</sup>) |
| --- | --- | --- | --- | --- |
| **Fum\*** | 0.94 | 1.066 | 0.212 | 0.90 |
| **Succ\*** | 0.82 | 1.219 | 0.494 | 0.90 |
| **Mal\*** | 8.96 | 0.112 | 0.012 | 0.94 |
| **aKG** | 1.86 | 0.538 | 0.021 | 0.92 |
| **GAP** | 0.22 | 4.568 | 3.573 | 0.76 |
| **Glc\*** | 3.84 | 0.261 | 0.485 | 0.96 |
| **Cit\*** | 260.96 | 0.004 | 0.008 | 0.89 |
| **iCit** | 6.65 | 0.150 | 0.001 | 0.99 |
| **2PG** | 1.94 | 0.515 | -0.005 | 0.93 |
| **DHAP** | 1.06 | 0.940 | 0.082 | 0.91 |
| **3PG** | 20.05 | 0.050 | 0.005 | 0.95 |
| **E4P** | 0.04 | 25.442 | 8.540 | 0.68 |
| **Rib5P** | 3.04 | 0.329 | 0.001 | 0.92 |
| **Ribu5P** | 1.04 | 0.965 | 0.099 | 0.76 |
| **Xyl5P** | 1.31 | 0.763 | 0.012 | 0.97 |
| **M6P** | 11.70 | 0.085 | 0.002 | 0.91 |
| **F6P** | 7.24 | 0.138 | -0.001 | 0.93 |
| **G6P** | 34.00 | 0.029 | 0.002 | 0.92 |
| **6PG** | 3.03 | 0.330 | -0.015 | 0.85 |
| **S7P** | 10.58 | 0.094 | -0.005 | 0.92 |
| **Trehalose** | 626.57 | 0.002 | 0.003 | 0.96 |

\* The concentrations of these compounds were calculated from injections measured in split (1:5) mode.

**Table 13.** Quantification of amino acid concentrations in the internal standard using calibration with commercial compounds. The reported concentration has been corrected for the addition of 10 µL internal standard to the calibration sample. All samples were measured in duplicate.

| Compound | Concentration [µM] | Slope | Intercept | Linearity (R<sup>2</sup>) |
| --- | --- | --- | --- | --- |
| **Alanine** | 112.98 | 0.089 | -0.035 | 0.985 |
| **Glycine** | 27.19 | 0.368 | 0.195 | 0.981 |
| **Valine** | 116.73 | 0.086 | -0.006 | 0.988 |
| **Leucine** | 17.76 | 0.563 | 0.004 | 0.990 |
| **iso-Leucine** | 33.49 | 0.299 | 0.011 | 0.989 |
| **Proline** | 6.11 | 1.637 | -0.041 | 0.982 |
| **Serine\*** | 76.28 | 0.131 | 0.001 | 0.999 |
| **Threonine** | 61.46 | 0.163 | -0.012 | 0.987 |
| **Methionine\*** | 1.27 | 7.886 | 0.143 | 0.981 |
| **Aspartate** | 266.64 | 0.038 | -0.012 | 0.985 |
| **Phenylalanine\*** | 24.81 | 0.403 | 0.001 | 0.987 |
| **Glutamate** | 1967.34 | 0.005 | 0.007 | 0.984 |
| **Lysine** | 415.89 | 0.024 | 0.003 | 0.983 |
| **Asparagine** | 125.63 | 0.080 | -0.005 | 0.982 |
| **Glutamine** | 901.14 | 0.011 | 0.004 | 0.986 |
| **Tyrosine\*** | 61.21 | 0.163 | 0.022 | 0.991 |
| **Histidine** | 374.45 | 0.027 | -0.002 | 0.979 |
| **Tryptophane\*** | 22.27 | 0.449 | 0.024 | 0.991 |

\* The concentrations of these compounds were calculated from splitless injections.

For the glycolytic intermediates, the metabolites GAP, E4P, Ribu5P, and 6PG could be detected by GC-MS analysis, however, the concentrations were too low for accurate quantification. These sugar phosphates and molecules with a high molecular mass are often more difficult to measure since they are more sensitive to interruptions in the lyophilization process, exposure to sunlight, and matrix effects (Niedenführ et al., 2016).

## Basic Protocol 8: Quantification of metabolites in yeast samples

Once the internal standard has been characterised as described in Basic Protocols 6 and 7, it is ready to be used for quantification of intracellular metabolites in cell extracts. This protocol describes how to perform cultures for quantification of intracellular metabolites using internal standards.

### Materials

*S. cerevisiae* strain IMX2600 (van den Broek et al., 2024) or other strain (1 mL in 30% v/v glycerol)

Synthetic mineral medium supplemented with 2% glucose and vitamin solution (Verduyn et al., 1992) or other growth medium of interest

Methanol (HPLC-grade >99.9%) (-40 °C), 100% and 83% (v/v) 75% (v/v) ethanol solution

Internal standard (^13^C yeast cell extract)

Round bottom 500 ml shake flasks

Innova 4000 incubator shaker (Eppendorf) at 30 °C

Jenway 7200 scanning spectrophotometer (Cole-Parmer Inc, Chicago, USA)

Cryostat with cryofluid (70% v/v ethylene glycol), set at -40 °C Freezer, set at -40 °C

### 15 mL Greiner tubes filled with 5 mL 100% methanol

Scale

Pipetboy and sterile 2 mL pipettes

Vortex

Centrifuge (tabletop, -19 °C) – rotor for 15 mL tubes

Bottle-top dispensers (5 mL and 20 mL capacity)

Waste bucket for methanol waste (plastic and liquid)

Two water baths (75 °C, and 95 °C)

Regular marbles (14 mm diameter)

Benchtop centrifugal vacuum concentrator (SpeedVac), e.g. Labconco, Eppendorf or equivalent, with vacuum pump and cold trap, capable of evaporating ethanol and equipped with a rotor for 15 mL conical tubes

Syringes

0.2 μm syringe filters (PVDF)

Nitrile gloves

### Protocol

1. Grow a strain of interest in a shake flask or bioreactor. Bioreactors are preferred for intracellular metabolite analysis as parameters (pH, gases, nutrients) can be tightly controlled, including (limited) nutrient supply. Additionally, bioreactors, like the ones used in the present study, can be equipped with sampling ports enabling ultrafast sampling within milliseconds (*Figure 2*), which minimizes metabolic changes during sampling. However, as shown in this protocol, sampling from shake flask cultures is also possible, provided that gas and nutrients are not in limited supply for cells during sampling.
2. Weigh 5 mL of methanol in a 15 mL Greiner tube and record the exact weight. Store at -40 °C.
3. Record the optical density of the culture.
4. Once at the desired optical density, sample 1.2 mL of culture into 5 mL of -40 °C methanol as quickly as possible using a pipetboy with 2 mL pipette. To be above the detection limit of the mass spectrometer, the sampling volume must be adjusted to the cell density of the culture and the expected intracellular metabolite levels. The sampling volume may be decreased or increased as long as the quenching ratio of 1:5 (culture: methanol) is maintained.
5. Immediately vortex the sample.
6. Weigh the broth-methanol sample and record the exact weight and keep the sample on dry-ice or in a -40 °C cryostat. As mentioned previously, to arrest all metabolic activity the samples need to be kept cold until metabolite extraction using boiling ethanol.
7. Perform the sample processing and metabolite extraction according to Basic Protocol 2 steps 7 to 34 with the following adjustments.

a. In step 9 of Basic Protocol 2, add 5 mL 83% (v/v) methanol (-40 °C) instead of 20 mL.
b. Skip step 12 of Basic Protocol 2 because Basic Protocol 8 only works with 15 mL Greiner tubes and therefore the rotor does not need to be changed.
c. Skip step 13 of Basic Protocol 2 because the second washing step will use 5 mL 83% methanol again.
d. In step 15 of Basic Protocol 2, add 5 mL 83% (v/v) methanol (-40 °C) instead of 3 mL.
e. Skip steps 17 to 19 of Basic Protocol 2 because the transfer from 50 to 15 mL tubes is not needed.
f. After step 21 of Basic Protocol 2, add internal standard to the cell suspension which is mandatory for quantification of intracellular metabolites. Additionally, ^13^C-NA was spiked at a final concentration of 100 µM since this compound is not present in the internal standard.
g. In step 25 of Basic Protocol 2, boiling for 3 minutes is sufficient.
h. In step 29 of Basic Protocol 2, we have used a MilliQ volume of 600 µL to resuspend the pellets and found that this resulted in a measurable metabolite concentration for *S. cerevisiae* cultures grown on chemically defined medium with glucose as sole carbon source during exponential growth (OD_660_ 2-8) *In this protocol, 100 µL of internal standards prepared in Basic Protocol 5 are spiked to each tube. The volume of internal standard to add is optimized to meet the detection limit of the mass spectrometer. While the volume of internal standard used should be minimized to reduce cost, ^13^C-labelled metabolites should be clearly detectable by the mass spectrometer*.
8. Perform LC- or GC-MS analysis following Basic Protocol 6 or 7.
9. See the Understanding of Results section to learn how to calculate the intracellular metabolite concentrations based on mass spectrometry output.

### Reagents and solutions

#### Antifoam C (1/10) solution

Add 10 mL of antifoam C emulsion (shake well before use) to a bottle. Add 90 mL of demi-water and shake well.

Sterilise at 121 °C for 20 minutes.

#### Vitamins solution (1 L)

Calibrate the pH probe using pH 4 and pH 7 buffers.

Take a Schott bottle (1 L) and add a stirrer bar.

Fill the Schott bottle with 1000 mL of demi-water and mark the volume.

Remove approximately 200 mL demi-water from the bottle.

Dissolve 0.05 grams of D-biotin in 10 mL of 0.1 M NaOH in a Greiner tube.

Add the biotin solution to the 800 mL MilliQ present in the Schott bottle.

Put the bottle on a magnetic stirrer plate.

Adjust the pH to 6.5 with HCl (1 M).

Add 1 gram of thiamine and dissolve properly.

Adjust the pH to 6.5 with NaOH (1 M). Adjust final volume to 1 L.

Filter sterilise the vitamin solution and store in the fridge (up to 3 months).

#### Trace elements solution (5 L)

Calibrate the pH probe using pH 4 and pH 7 buffers.

Take a 5 L flask and add a stirrer bar.

Fill the flask with demi-water and mark the 5 L level.

Remove approximately 1.25 L of MilliQ water.

Put the flask on a magnetic stirrer plate.

Dissolve 75 grams of EDTA and 22.5 grams of zinc sulfate heptahydrate (ZnSO_4_ .7H_2_O).

Adjust the pH to 6.0 with NaOH (1 M).

Add all the following compounds one by one in the specified order and correct the pH to 6.5 after each

addition.

*NOTE*: Wait until each compound has completely dissolved before adding the next compound.

- 4.2 grams of manganese (II) chloride ·2H_2_O
- 1.5 grams of cobalt (II) chloride · 6H_2_O (**toxic**)
- 1.5 grams of copper (II) sulfate · 5H_2_O
- 2 grams of di-sodium molybdate · 2H_2_O
- 22.5 grams of calciumchloride · 2H_2_O
- 15 grams of iron sulfate · 7H_2_O
- 5 grams of boric acid
- 0.5 grams of potassium iodide

When all compounds have dissolved, adjust the pH to 4.0 with HCl (1 M). Add demi-water to adjust the volume to 5 L.

#### pH titration solution (4 M NH_4_OH) (200 mL)

*NOTE:* NH_4_OH has high vapour pressure and the gas is toxic. Always work in the fume hood and **do not** sterilise this solution.

Add 88 mL of MilliQ to a Schott bottle (250 mL).

Sterilise this bottle with MilliQ at 121 °C.

Sterilise the screwcap with two in/outlets separately in a bottle.

In a fume hood, add 112 mL of NH_4_OH.

Transfer the sterilised screwcap with three distributors to the base solution bottle.

#### Glucose solution (100 mL)

Take a Schott bottle (250 mL) and add a stirrer bar.

Fill the Schott bottle with 100 mL MilliQ and mark the volume. Remove approximately 20 mL of MilliQ.

Put the bottle on a magnetic stirrer plate.

Dissolve 6.81 grams of ^12^C-glucose monohydrate (TRIAL RUN) or 6.39 grams ^13^C-glucose anhydrous (REAL RUN).

*NOTE*: Anhydrous ^13^C-glucose might dissolve poorly. To speed up its dissolving. put the solution in a 37 °C water bath.

Adjust the volume to 100 mL.

Sterilize the glucose solution at 110 °C.

#### Batch salt base (350 mL)

Calibrate the pH probe using pH 4 and pH 7 buffers.

Take a Schott bottle (500 mL) and add a stirrer bar.

Fill the Schott bottle with 350 mL of demi-water and mark the volume.

Remove approximately 100 mL demi-water from the bottle.

Put the bottle on a magnetic stirrer plate.

Dissolve the following compounds:

- 0.5 grams of ammonium sulfate (NH_4_)_2_SO_4_
- 1.5 grams of ammonium dihydrogen phosphate NH_4_H_2_PO_4_
- 1.76 grams of potassium dihydrogen phosphate KH_2_PO_4_
- 0.5 grams of magnesium sulfate heptahydrate MgSO_4_ · 7H_2_O
- 1 mL of trace elements solution
- 0.75 mL of anti-foam C (1/10) solution

Adjust the pH to 5 with KOH (2 M).

Adjust the volume to 350 mL.

Sterilize at 121 °C.

#### Fed-Batch medium (640 mL)

*NOTE:* It is recommended to prepare and filter sterilise the fed-batch medium on the day it will be used.

Calibrate the pH probe using pH 4 and pH 7 buffers.

Add 1 mL of anti-foam C (1/10) solution to a Schott bottle (1 L)

Sterilize the bottle at 121 °C.

Fill a non-sterile Schott bottle (1 L) with 640 mL of demi-water and mark the volume.

Remove approximately 100 mL demi-water from the bottle.

Put the bottle on a magnetic stirrer plate and add a stirrer bar.

Dissolve the following compounds:

- Glucos**e:**

`o` **TRIAL RUN**: 74.52 grams of ^12^C-glucose monohydrate
`o` **REAL RUN**: 70 grams of ^13^C-glucose anhydrous
- 0.64 grams of ammonium sulfate (NH_4_)_2_SO_4_
- 1.92 grams of ammonium dihydrogen phosphate NH_4_H_2_PO_4_
- 2.24 grams of potassium dihydrogen phosphate KH_2_PO_4_
- 0.64 grams of magnesium sulfate heptahydrate MgSO_4_ · 7H_2_O
- 3.2 mL of trace elements solution
- 3.2 mL of vitamin solution

Adjust the pH to 5 with KOH (2 M).

Remove the stirrer bar.

Adjust the volume to 640 mL.

Filter sterilise using a bottle-top filter and the vacuum pump into the sterilized bottle containing anti-foam C solution.

#### Mobile phase A & Weak wash: 50 mM ammonium bicarbonate in MS-grade H_2_O (1 L)

Weigh 3.953 g of ammonium bicarbonate and add to 1 L of Optima LC/MS grade H_2_O.

#### Strong wash & Seal wash: 10% acetonitrile, 10 mM ammonium bicarbonate in H_2_O (1 L)

Weigh 0.791 g of ammonium bicarbonate and to a 1 L flask.

Add 900 mL Optima LC/MS grade H_2_O to the 1 L flask.

Add 100 mL Optima LC/MS grade acetonitrile to the 1 L flask.

#### 10 mM ammonium bicarbonate in 60% acetonitrile (50 mL)

Weigh 39.53 mg ammonium bicarbonate and add to 30 mL Optima LC/MS-grade acetonitrile and 20 mL Optima LC/MS-grade water in a 50 mL Falcon tube.

#### Critical parameters and troubleshooting

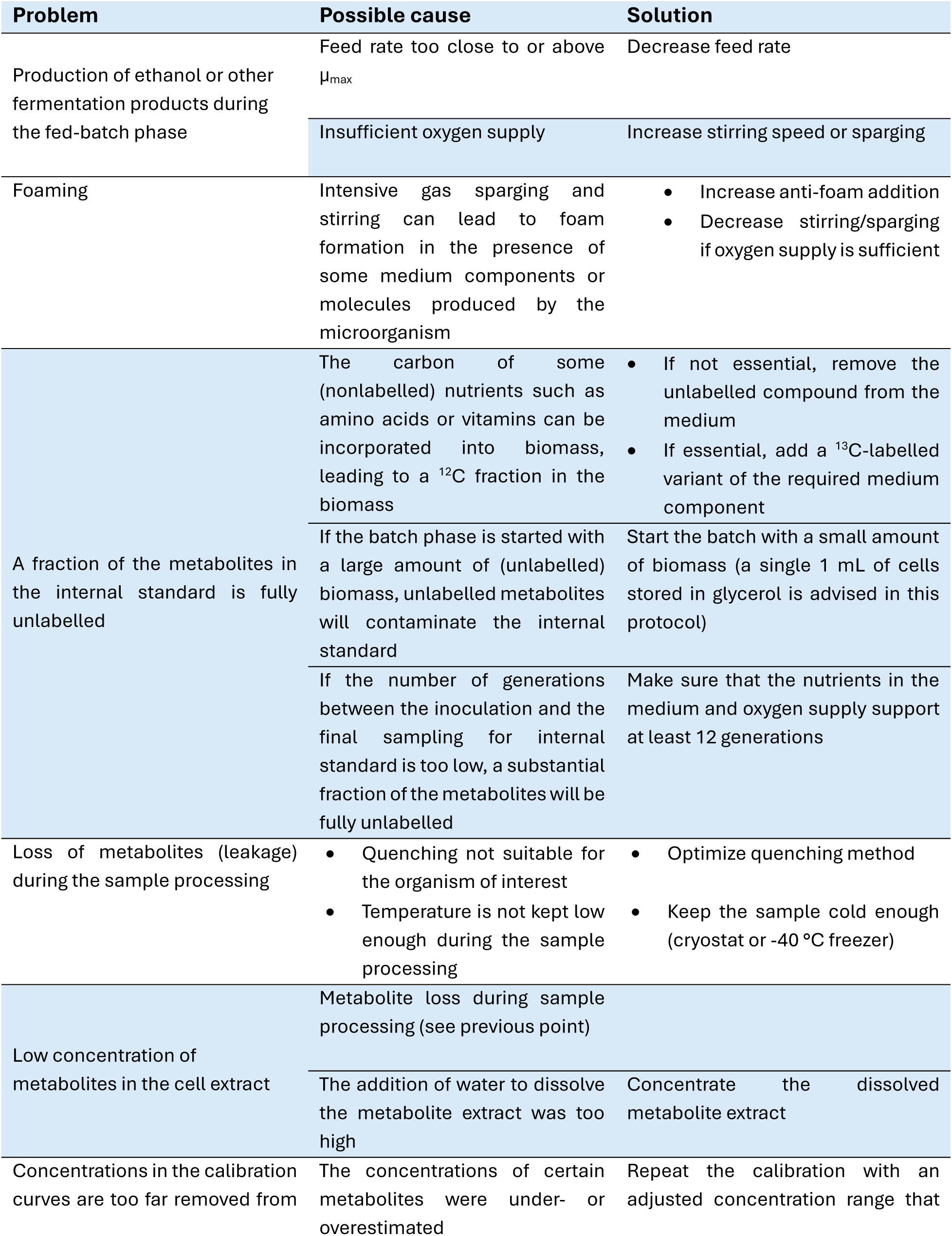
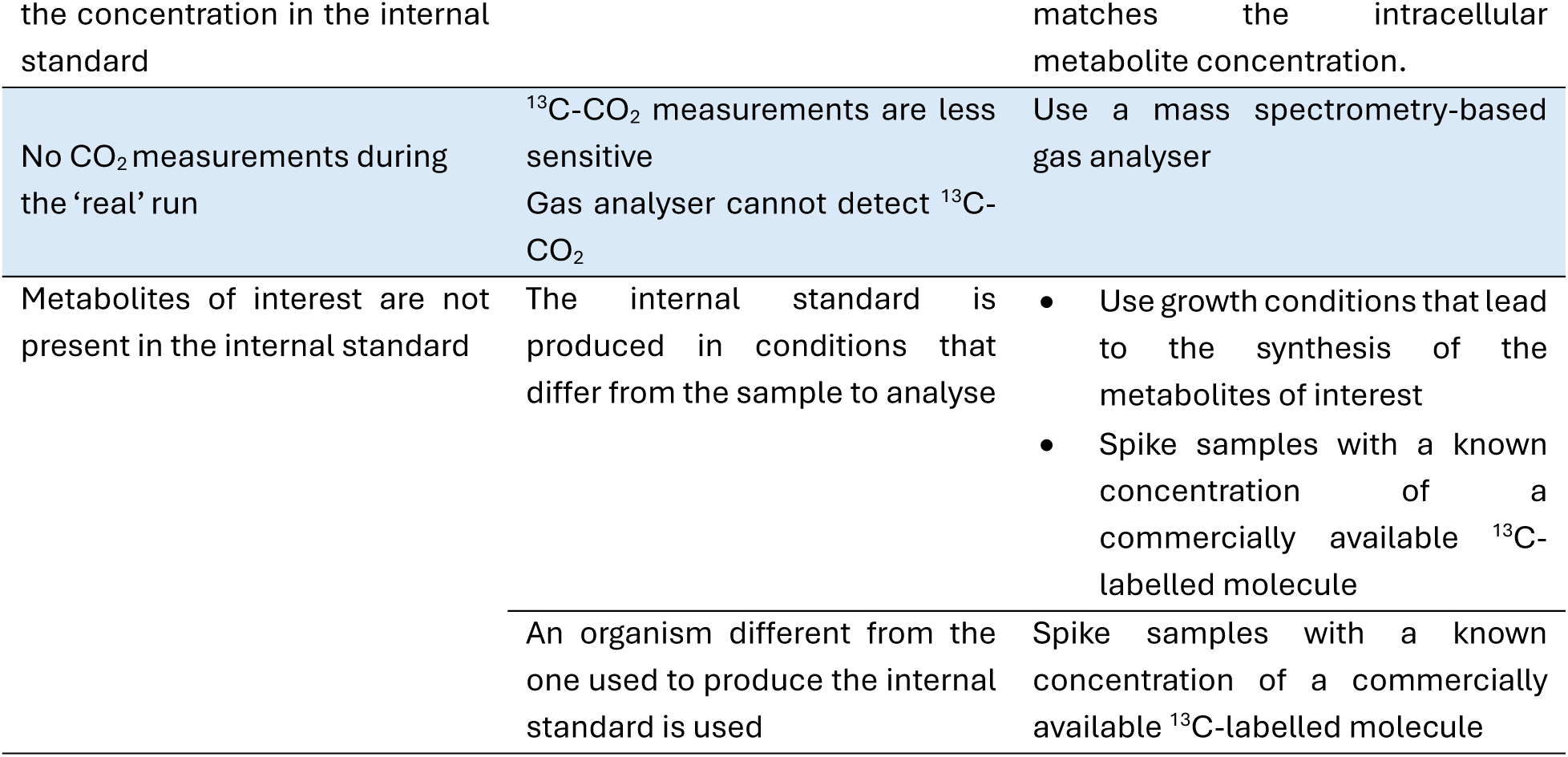

#### Understanding results

This section explains how to obtain intracellular metabolite concentrations by processing the data generated in Basic Protocol 8. The calculations can be performed using the Excel sheet in Supplementary File 3. For more detailed information on these calculations, please refer to Wahl et al. (2014).

1. Visualize and manually perform the quality control of the mass spectrometry data using XCalibur or other similar software.

2. For each metabolite, extract the ^12^C and ^13^C peak intensities. We used a Matlab script to extract the peak intensities, and the Matlab script can be provided upon request.

3. Calculate the ^12^C/^13^C ratio using equation 6 or use the spreadsheet ‘Ratio’ in Supplementary File 3.

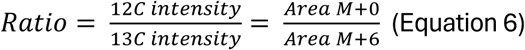

4. Enter the slope, intercept and quality of fit of the calibration curves in the spreadsheet ‘Calibration Curves’ in Supplementary File 3.

5. Use the ^12^C/^13^C ratio to calculate the measured amount of corresponding metabolite using the calibration equation (Spreadsheet ‘Metabolite Amount LCMS Sample’ in Supplementary File 3). This step includes a correction for final amount of water that the metabolite extract was dissolved in prior to LC or GC-MS analysis (20 µL).

6. Enter the sampled culture volume and corresponding optical density at 660 nm (OD₆₆₀) in the spreadsheet “Growth Curves” in Supplementary File 3. The spreadsheet will convert the optical density into dry biomass (gDW) concentration at sampling time using a pre-established OD_660_–dry-weight correlation of yeast.

7. The spreadsheet ‘Metabolite Amount umol per gDW’ in Supplementary File 3 multiplies the sampling volume by the dry-weight concentration (g/L) to obtain the weight of dried biomass used at the time of sampling for each sample. It will then correct the LC/GC–MS–quantified metabolite amount with this dry weight to generate the metabolite concentration expressed in µmol metabolite per gram biomass dry weight.

8. Finally, bring the metabolite concentration to µM by correcting with the intracellular volume per gram of dry biomass as described in the spreadsheet ‘Metabolite Amount uM’ in Supplementary File 3. In yeast an intracellular volume of 1.7 mL per gram of dry weight is typically used. Figure 8 represents the intracellular concentration of a subset of metabolites in nucleotides metabolism calculated in Supplementary File 3 using custom-made internal standards. Samples were taken as described in Basic Protocol 8 from cultures in shake-flasks.

**Figure 8.**
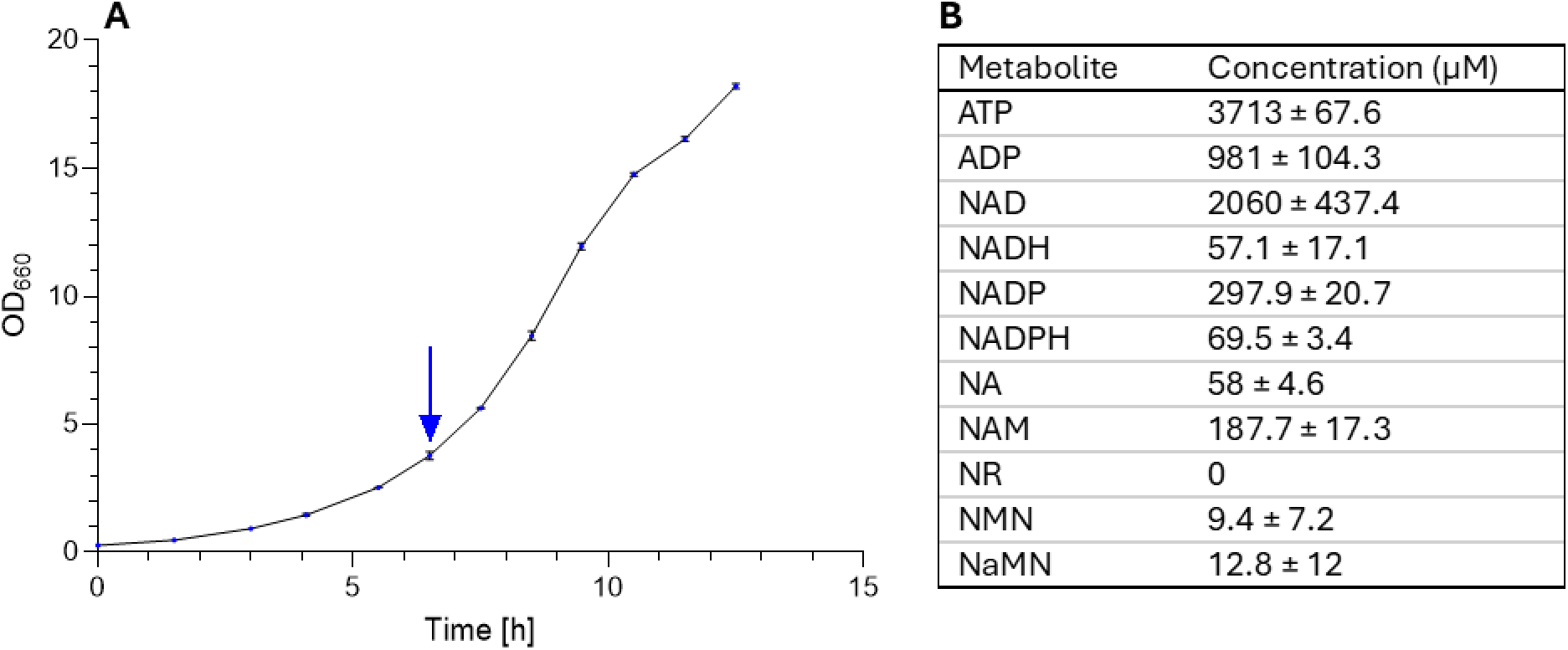
Intracellular metabolite concentrations in S. cerevisiae cultures measured using custom-made internal standards. S. cerevisiae IMX2600 (van den Broek et al., 2024) was grown in aerobic shake-flask chemically defined medium with glucose as sole carbon source. (A): biomass profile, (B): intracellular metabolites concentrations at time point 6.5 hours. Data represent the average and standard deviation of three independent culture replicates.

*Nicotinamide Riboside is an unstable compound that rapidly degrades during sample processing. It can be detected, but the measured concentrations are unreliable*.

*The measured NADPH/NADP⁺ ratio was outside of the expected range (expected ca. 1; see e.g. Zhang et al. (2015)), although both compounds providing a strong, quantifiable signal. Despite careful operation of the column (properly grounded, thoroughly cleaned and stored) and the prevention of redox-active additives (e.g., formic acid), the conductive surface of the porous graphitic carbon (PGC) stationary phase might have affected the detection of certain redox-sensitive analytes such as the NADP(H) couple (Bapiro et al., 2016; Melmer et al., 2010; Pabst et al., 2010; Törnkvist et al., 2004; van Ede et al., 2026)*.

#### Time considerations

As shown in Figure 1, the in-house production and characterization of internal standards require the following time allocation for *S. cerevisiae*:

- Ca. 5 days for the batch phase (1 day of medium preparation, 1 day for autoclaving, 3 days (∼ 66 hours) of fermentation).
- Ca. 2 days for the fed-batch phase of the fermentation (∼ 55 hours).
- 1 day for the sampling.
- 4 days for the extraction of metabolites, however the duration depends on the number of samples to process.

`o` The quenching of the samples and further processing until after the boiling ethanol step for metabolite extraction (step 25 in Basic Protocol 2) is time sensitive as metabolites might be unstable if the samples are not strictly kept at low temperature close to 0°C.
`o` The time required for ethanol evaporation depends on the capacity of the vacuum evaporator. Considering the standard capacity of a vacuum evaporator and the number of samples to process, this step will take multiple days.
- Ca. 4 days for quantification (1 day for preparation of calibration curve samples, 1 day for running all the samples, and 2 days for data analysis)

Note that the calibration curves and the amount of internal standard added to each sample might have to be optimized. It is essential to have a calibration curve in the correct range – around a ^12^C/^13^C ratio of 1 – to prevent under- or overestimation of metabolite concentrations. As a results, consider that this step might take two or three iterations when the methods is used for the first time.

The above-indicated time investment is for one successful run to produce, extract, and characterize the internal standard. Please consider that you will have to go through this cycle at least two times to perform a trial run to test the yield of biomass on substrate and to identify whether the metabolites of interest are present in the cell extract. The trial run might have to be executed multiple times.

## Acknowledgements

We are grateful for Marijke Luttik, Yanfang Wang, Marieke Warmerdam, Tobias Fecker, Bas Michels, and Martijn Sandelowsky experimental support for the sampling and processing of cell samples.

M.C.’s PhD project is funded by the Dutch Research Council (NWO) OCENW.XL21.XL21.007 grant. R.W.’s PhD project is funded by a Stevin Prize awarded to Jack Pronk by NWO. J.vE.’s PhD project is funded by the TU Delft Zero Emission Biotechnology program.

## Data availability statement

All acquired data are publicly accessible through the 4TU.ResearchData platform, DOI 10.4121/89ba9ee3-f072-4dda-97ba-683cce975a3b. The LC-MS data analysis MATLAB script is available upon request.

## Supporting Information

Supplementary File 1, Excel file, can be used to simulate the fed-batch feeding profile to estimate the final biomass concentration and to determine the sampling points.

Supplementary File 2, text file, describes how to calculate the fed-batch feeding profile. Supplementary File 3, Excel file, can be used to calculate metabolite concentrations from mass spectrometry data.

Supplementary File 4, Excel file, shows how to calculate the amount of ^12^C contamination in the internal standard.

