## Supplementary File 2 for "A protocol for lab-scale production of ^13^C yeast extract as internal standard for metabolomics and quantification of intracellular metabolites"

This readme file was generated on 2024-01-26 by Rosalie Wouters and Miriam Cammaert

 

GENERAL INFORMATION

Title of Dataset: Fed-batch calculations for 13C fermentation of yeast strain CEN.PK113-7D


Principal Investigator Information

Name: Prof. Pascale Daran-Lapujade

Institution: Delft University of Technology

Address: Van der Maasweg 9, 2629HZ, Delft



 

Author Information

Name: Rosalie Wouters; Miriam Cammaert

Institution: Delft University of Technology

Address: Van der Maasweg 9, 2629HZ, Delft




This file contains the theoretical background for the calculations performed in Excel file '20240126_Simulation_Fedbatch'
In part 1, the calculations behind the determination of the fedbatch feeding profile are explained -including the adjustments in feeding profile required after sampling.
In part 2, the practical aspects of the Excel file -i.e., how to use it properly- are discussed.  

#1. Determination fedbatch feeding profile

To model the feeding rate required for an exponentially growing culture with a set growth rate (mu), a couple of parameters are required. 
 
mu [h-1]: specific growth rate, set by feeding rate
ms [g_S g_X-1 h-1]: maintenance requirements, organism dependent
Y_X_S [g_X g_S-1]: yield of biomass on substrate, organism dependent
V_0 [L]: starting volume in the reactor
c_X_0 [g_X L-1]: starting biomass concentration in the reactor
c_S_in [g_S L-1]: substrate concentration in the fedbatch medium 

The feeding profile for the fedbatch is derived from the Herbert-Pirt relation (Eq. 1)

(a) q_S = mu/Y_X_S + q_P/Y_P_S + ms (Eq. 1)
	With:
	mu [h-1]; specific growth rate
	q_S [g_S g_X-1 h-1]: biomass specific substrate consumption rate
	q_P [g_P g_X-1 h-1]: biomass specific product formation rate
	Y_X_S [g_X g_S-1]: yield of biomass on substrate
	Y_P_S [g_P g_S-1]: yield of product on substrate
	ms [g_S g_X-1 h-1]: maintenance requirements

(b) Assume zero product formation in an aerobic, glucose-limited culture (q_P = 0)

(c) Combine (a) & (b)

    q_S = mu/Y_X_S + ms (Eq. 2)

(d) Define the total amount of biomass in the reactor at timepoint zero (M_X_0)
    
    M_X_0 = c_X_0*V_0 (Eq. 3)
        With:
	M_X_0 [g_X]: starting total amount of biomass in the reactor (t = 0)
	c_X_0 [g_X L-1]: starting biomass concentration in the reactor (t = 0)
	V_0 [L]: starting volume in the reactor (t = 0)


(e) Define the amount of biomass in the reactor as a function of time (M_X)

    M_X = M_X_0 * exp(mu*t) (Eq.4)
	With:
	M_X [g_X]: biomass at timepoint (t) in the reactor
	t [h]: timepoint 

(f) Define the total substrate consumption rate in the reactor (R_S)

    R_S = q_S * M_X (Eq. 5)
	With:
	R_S [g_S h-1]: total substrate consumption rate in the reactor


(g) Combine Eq. 2, Eq. 4 & Eq. 5

    R_S = (mu/Y_X_S + ms) * c_X_0 * V_0 * exp(mu*t) (Eq. 6)

(h) Define the volumetric substrate feedrate (F_S)

    F_S = R_S/c_S_in (Eq. 7)
	With: 
	F_S [L h-1]: volumetric substrate feedrate 
	c_S_in [g_S L-1]: substrate concentration in the fedbatch medium 

(i) Combine Eq. 6 & Eq. 7
	
    F_S = (mu/Y_X_S + ms) * ((c_X_0 * V_0)/c_S_in) * exp(mu*t) (Eq. 8)

(j) Simplify Eq. 8

    F_S = f * exp(mu*t) (Eq. 9)
	With: 
	f [L h-1]: (mu/Y_X/S + ms) * ((c_X_0 * V_0)/c_S_in)

(k) Eq.8 allows for determining the feed profile for the fedbatch phase, up until the first sample. 
    After taking the sample at timepoint 't' the feedrate needs to be adjusted, as both the total volume in the reactor (l) and the total amount of biomass (m) in the reactor change.

(l) The new volume (V) can be determined by the sum of feed addition and sample removal.
    NOTE: for simplification purposes, pH controle and antifoam addition are neglected in the total volume. 

    V = V_0 + ΔV_feed - ΔV_sample (Eq. 10)
	With 
	V [L]: volume at timepoint (t)
	ΔV_feed [L]: total change in volume at timepoint (t) due to feed addition
	ΔV_sample [L]: total change in volume at timepoint (t) due to sampling
    
    The change in volume due to feed addition (ΔV_feed) can be obtained by integration of Eq. 9

    ΔV_feed = ∫ Fs = (f/mu)*exp(mu*t) - (f/mu) (Eq. 11)

    The change in volume due to sampling (ΔV_sample) can be obtained by determining the volume of broth removed during sampling 
    (NOTE: in all our calculations, we assumed the density of water for our samples, and resultingly neglected the effect of the biomass on the sample density)
	 
    Allows for substitution in Eq. 10

    V = V_0 + (f/mu)*exp(mu*t) - (f/mu) - ΔV_sample (Eq. 12)

(m) The total amount of biomass in the reactor after sampling (M_X_#) can be determined based on the volume of the sample (ΔV_sample) as a fraction of the total volume (V).
    
    M_X_# = M_X * (1 - (ΔV_sample/V)) (Eq. 13)

    NOTE: Due to the discrepancy between exponential increase of the total volume in the reactor and the total amount of biomass in the reactor, it is important to calculate the total amount of biomass rather than the biomass concentration for determining the feeding profile. 

(n) The total amount of biomass in the reactor after sampling (M_X_#) can be considered as a new starting time point (t = 0) and as such, the calculations for determining the exponential feedrate can be repeated starting from (d), assuming the new total biomass amount and total volume adjusted for sampling. 


#2. How to use the Excel file to calculate the feedrate adjustments 

(a) Open Excel file '20240126_Simulation_Fedbatch'

(b) Open the 'FEDBATCH SIM' tab

(c) Fill in the required parameters, highlighted in yellow: 
     
    mu [h-1]: specific growth rate, set by feeding rate
    Y_X/S [g_X g_S-1]: yield of biomass on substrate, organism dependent
    ms [g_S g_X-1 h-1]: maintenance requirements, organism dependent
    V_0 [L]: starting volume in the reactor
    c_S_in [g_S L-1]: substrate concentration in the fedbatch medium 
    c_X_0 [g_X L-1]: starting biomass concentration in the reactor (at the start of the fedbatch)

(d) Fill in the approximated fed-batch time and check the total amount of fed-batch medium added. In our case, we aimed for an addition of approximately 0.490 L, as this leaves enough medium during the sampling day based on a total of nine samples over a period of ten hours. 

(e) Open the 'FEDBATCH SAMP' tab. The yellow spaces, again, indicate values that have to be filled out. The parameters as defined on the other tab (see (c)) do not have to be filled out again, but are already defined. 
    Sample 1 reflects the first sample taken after the fedbatch phase at the time point defined in the previous step (d). 
    
(f) Fill in the weight of the sampling container ('Beaker [g]') and the weight of the sampling container filled with culture ('Beaker + culture [g]') to determine the amount of volume removed from the reactor ('Culture [g]'; again, assume density of water, i.e., 1 kg L-1)
    After sampling, a new fedbatch profile has to be calculated as the amount of biomass in the reactor has changed. As such, the sampling time point is considered as a new time point zero (t = 0). 
    This resetting of the time is reflected in a new row, wherein the adjusted feeding rate constant ('f') is calculated. 
    This adjusted feeding rate constant ('f') should be incorporated into the feeding rate and it allows for the calculation of the total volume at the following sampling point

(g) For all the folllowing sampling steps, step (f) can be repeated. 

(h) For all the sampling steps, ensure that the biomass concentration in the reactor before and after sampling is the same, as this indicates that the calculations of the biomass removal during sampling are correct. Additionally, it is recommended to check the theoretical amount of medium added based on the sampling schedule before starting the experiment. 
